# Cell Cluster Geometry and Fluidity Control the Transition from Single-Cell Chemorepulsion to Collective Chemotaxis

**DOI:** 10.64898/2026.07.04.736449

**Authors:** Monika Sanoria, Gema Malet-Engra, Giorgio Scita, Nir Gov, Ajay Gopinathan

## Abstract

Directed migration along chemical gradients controls immune surveillance, development, and cancer invasion. However, the same chemical cue can produce different responses depending on its concentration and whether cells move alone or in groups. For example, in steep gradients, isolated malignant lymphocyte cells migrate away from the chemoattractant source, whereas clusters of the same cells continue to migrate toward it. Here, combining computational modeling and experimental observations, we show that this reversal is governed by coupled mechanisms acting across molecular, cellular, and collective scales. At the single-cell level, our model predicts that receptor endocytosis generates a feedback that produces a nonmonotonic surface receptor density with increasing chemoattractant concentration. Above a critical concentration that depends on the cell’s volume-to-sensing-area ratio, receptor depletion reverses cell polarity and drives chemorepulsion. However, in clusters, cell-cell contacts reduce the membrane area exposed to ligand, increasing the volume-to-sensing-area ratio, thus increasing the critical concentration and preserving chemotaxis. An agent-based model incorporating these mechanisms quantitatively reproduces the sign reversal of the migration index across gradient steepness and cluster size. We show that collective rearrangements further stabilize chemoattraction with exchanges between the cluster rim and core helping remove chemorepulsive cells from the leading edge, keeping their fraction below the threshold required to reverse cluster migration. The model further predicts, and experiments confirm, that increasing ambient ligand concentration while keeping the gradient fixed reduces cluster chemoattraction. Our results identify receptor trafficking, cell geometry, and cluster fluidity as physical determinants of collective directional decision-making, with implications for immune cell homing, tissue morphogenesis, and cancer dissemination.

---

Directed migration along chemical gradients is fundamental to many physiological processes including immune surveillance and inflammatory responses [1, 2], cancer dissemination [3] and development [4]. Typically, a chemical (chemokine) spatial gradient biases cytoskeletal dynamics through asymmetric receptor activation, producing positive taxis in most systems [5, 6]. Chemorepulsion, defined as directed migration *away* from a signal, is less understood mechanistically, yet occurs for several chemokine-receptor pairs in physiologically relevant ranges [7–10] This complexity increases in collective migration, where cluster-level properties can produce novel emergent behavior [11–22] as seen in galvanotaxis [23] and in chemotaxis-driven cluster shape instabilities [24]. In many types of cancer, tumor clusters are the primary metastatic unit, colonizing distant sites more efficiently than single cells, actively assembling into heteroclonal aggregates and adopting collective migration modes [25–30].

One example is chronic lymphocytic leukemia (CLL),which is a B-cell malignancy in which tumor cells accumulate in lymph nodes through CCR7-receptor mediated homing to CCL19 chemical gradients[4]. Malet-Engra et al. [31] studied JVM3, a CLL-derived B-cell line, in microfluidic CCL19 gradients, and showed that, in shallow gradients, both single cells and clusters perform chemotaxis toward the source [31]. In steep gradients, however, their responses diverge. Single cells persistently migrate *away* from the source (chemorepulsion),while clusters of ≥ 20 cells persistently migrate *toward* the source (chemotaxis) (Fig. 1A–C).The same phenomenon is observed for primary CLL cells and for Jurkat T cells in steep CXCL12 gradients[31], suggesting a general phenomenon of collective lymphoid migration. It is clear that receptor trafficking is central to directional polarity [32–34], but the mechanism linking receptor dynamics to chemotactic response switching in single cells and how cell-cluster geometry can flip the sign of a chemorepulsive response remain open questions.

**FIG. 1.**
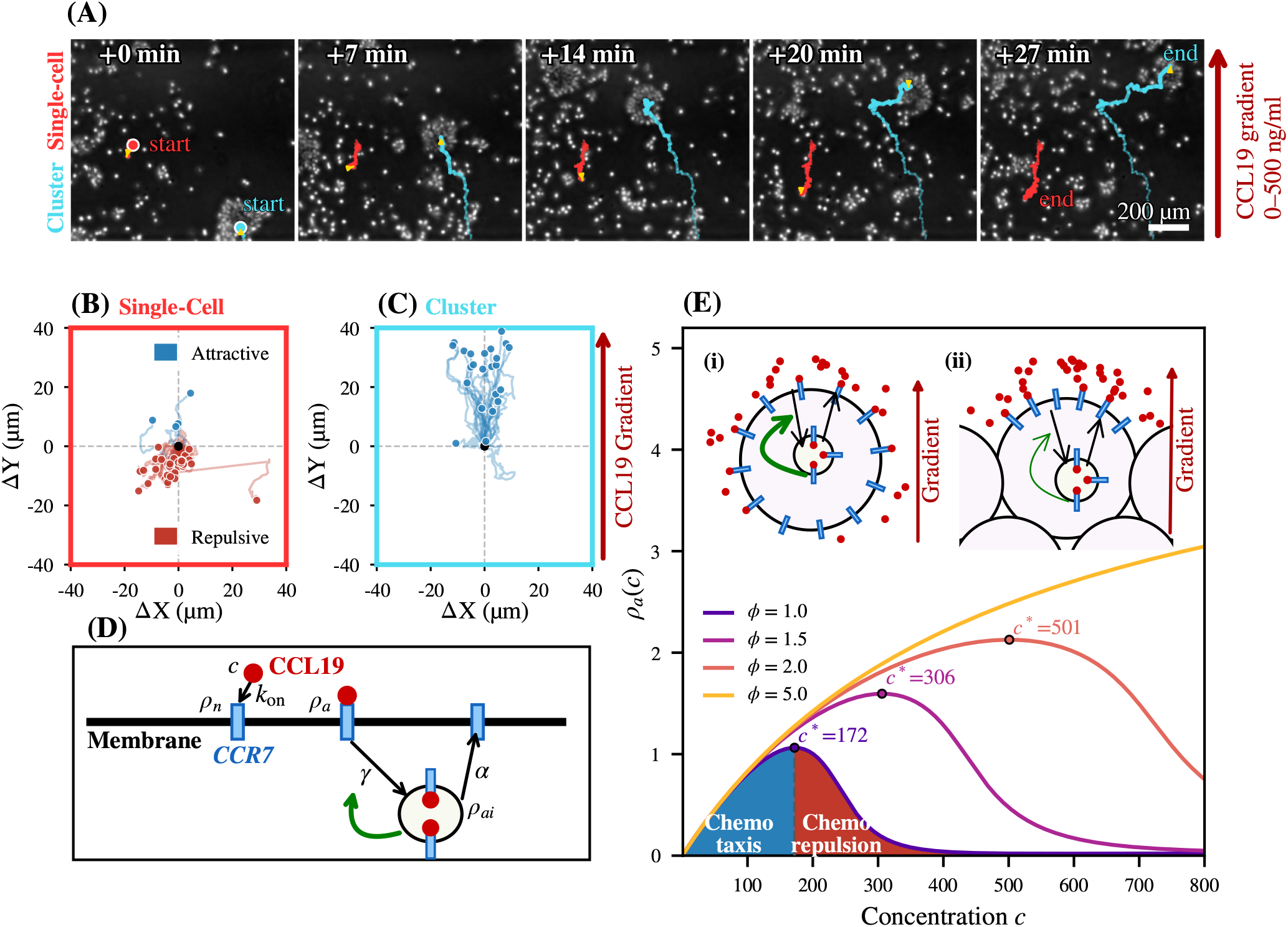
Single cell chemorepulsion, cluster chemoattraction and the receptor trafficking model. **(A)** Time-lapse microscopy of JVM3 (CLL) cells migrating under a 0–500 ng/mL CCL19 gradient (*t* = 0–27 min). A single cell (red trajectory) migrates away from the CCL19 source (chemorepulsion), while a cell cluster (cyan trajectory) migrates toward the source (chemoattraction). Gradient direction is indicated by the red arrow. **(B)** Displacement tracks of *n* = 43 single cells; majority show negative Δ*Y* (FMI = − 0.65). **(C)** Displacement tracks of *n* = 23 clusters; all positive Δ*Y* (FMI = +0.49). This sign reversal is the chemorepulsion-to-chemotaxis switch. **(D)** Schematic of the proposed model of the CCR7 trafficking circuit: 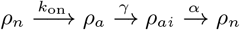 Accumulated *ρ*_*ai*_ triggers intracellular feedback (potentially ERK-mediated) that accelerates the endocytosis rate *γ* (green arrow), driving nonlinear receptor depletion above *c*. **(E)** Steady-state activated receptor density *ρ*_*a*_(*c*) for *ϕ* = 1.0, 1.5, 2.0, 5.0, using Eqs. (1–4). At *ϕ* = 1 (isolated single cell), *ρ*_*a*_ peaks at *c*\* ≈ 172 then falls; producing chemorepulsion; the blue- and red-shaded regions under the *ϕ* = 1.0 curve, separated by the dashed line at *c*\*, indicate the chemoattraction (*c < c*\*) and chemorepulsion (*c > c*\*) regimes, respectively; the feedback-ON state is illustrated in schematic (i). Increasing *ϕ* shifts *c*\* to 306 (*ϕ* = 1.5), 501 (*ϕ* = 2.0), and beyond the physiological gradient window (*ϕ* = 5.0), as *ϕ*-mediated receptor dilution progressively weakens the feedback (schematic (ii)). Parameters used *k*_on_ = 0.001, *γ*_0_ = 10.0, *α* = 8.0, *ε* = 200, *ρ*\* = 3.0, *σ*_*ρ*_ = 0.5, *ρ*_total_ = 10 (full list in Table I).

Prior work has explored aspects of this problem. At the collective scale, CCR7 acts as a chemokine-consuming sink in dendritic cells, with ligand internalization reshaping extracellular gradients to sustain long-range collective migration [31, 35]. This form of self-generated gradient guidance is well established across different biological systems [36, 37], but this extracellular mechanism does not address the intracellular switch that drives single-cell chemorepulsion. Recent work [38] has shown that a model where bound receptors drive a nonlinear feed-forward loop of intracellular signaling molecules can reproduce the single-cell chemorepulsion switch in JVM3 cells but it does not address the suppression of the switch in clusters.

Here we show that three nested mechanisms, operating at the molecular, cellular, and collective levels, jointly explain the experimental observations. At the *molecular* level, intracellular positive feedback on CCR7 endocytosis depletes surface receptors above a critical concentration, reversing cell polarity and driving single-cell chemorepulsion. At the *cellular* level, cell-cell contacts raise the volume-to-exposed-area ratio, diluting intracellular receptor feedback and shifting the critical concentration for chemorepulsion beyond the physiological gradient window, abolishing chemorepulsion in cell clusters. We implement these mechanisms in an agent-based model of cells migrating in two-dimensions, where stochastic polarity switching, calibrated from CCR7 kinetic data, quantitatively reproduces experimental observations. Our model reveals that, at the *collective* level, cluster fluidity continuously redistributes repulsive cells away from the leading edge through rim-core exchange, preventing their accumulation. Our results identify and establish receptor trafficking feedback, geometric volume-to-exposed-area and cluster fluidity as three critical determinants of chemotaxis and chemorepulsion in single cells and cell clusters.

## MODEL

We combine two complementary modeling frameworks: (i) an analytical model for receptor dynamics that predicts the critical concentration(*c*\*) above which endocytosis-driven receptor depletion reverses the polarity of isolated cells as a function of cell geometry, and an agent-based model (ABM) that simulates collective cluster migration and emergent rim-to-core cell exchange dynamics.

### (i) Analytical Receptor Dynamics Model

#### In Vivo Receptor Dynamics

The receptor CCR7, is a canonical G-protein-coupled chemokine receptor expressed on B lymphocytes and naïve T cells, essential for lymphocyte homing [4]. Its ligand-specific internalization kinetics make it susceptible to concentration-dependent trafficking feedback, making receptor endocytosis a candidate molecular switch for celll polarity. Specifically, CCL19 binding recruits *β*-arrestin-2 (an effect absent for the related ligand CCL21) and drives CCR7 internalisation via dynamin-dependent clathrin-coated pits. While CCR7 recycles back to the surface, CCL19 is degraded intracellularly [39, 40]. This recycling occurs via a ubiquitylation-dependent route [41], and internalized CCR7 continues to signal from endomembranes [42], suggesting that receptor trafficking can contribute non-trivially to cell polarity and migration direction [32–34].

Three observations from Malet-Engra et al. [31] implicate receptor trafficking in the chemorepulsion mechanism. First, dynamin inhibition converts single-cell chemorepulsion to chemoattraction, showing that endocytosis is required for the repulsive state. Second, leader cells at the cluster front show fluctuating CCR7 surface levels with frequent rim-core exchanges [31], suggesting higher chemorepulsive switching. Third, CCR7 surface reduction in leaders is consistent with a concentration-dependent positive feedback in which *β*-arrestin-2 amplifies endocytosis via two parallel pathways, an ERK pathway that increases clathrin-coated pit initiation [43–45] and a Src pathway that accelerates dynamin-mediated vesicle scission [46, 47]. Crucially, this ERK/FCHSD2 effect is cancer-cell selective [45], making it directly applicable to JVM3 cells, a malignant B-cell cancer line.

##### Mathematical Model of Receptor Dynamics

Based on the dynamics described above, we develop a coarse-grained model of CCR7 receptor trafficking capturing ligand binding, endocytosis, recycling, and an intracellular feedback signal (potentially ERK- and/or Src-mediated [39, 43–48]) that depends nonlinearly on the internalized receptor pool *ρ*_ai_ (model schematic in Fig. 1(D)). This feedback enhances the endocytic rate *γ*, enabling high ligand concentrations to drive a positive-feedback loop that can deplete surface receptors.

At the single-cell level, we track three CCR7 receptor pools: native (unbound) surface receptors *ρ*_n_, ligand-bound activated surface receptors *ρ*_a_, and internalized endosomal receptors *ρ*_ai_ (depicted in Fig. 1(D)). The non-dimensional equations for a cell of exposed membrane area *A*_exposed_ and volume *V*, with lengths rescaled by cell size *r*, are:

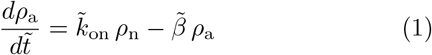

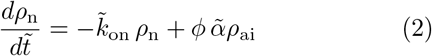

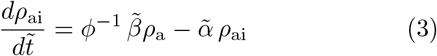

Here time is rescaled by the basal endocytosis rate 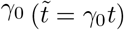, giving dimensionless parameters 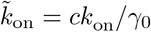 (ligand binding rate), 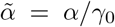 (recycling rate), and 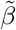 (feedback-modulated endocytosis; see Eq. 4), where *c* is the local chemokine concentration. Receptor number is conserved: *A*_exposed_*ρ*_n_ + *A*_exposed_*ρ*_a_ + *V ρ*_ai_ = *ρ*_total_ = *const*. Cell geometry enters through the volume-to-exposed-area ratio, *ϕ* ≡ *V/A*_exposed_. For a spherical cell of radius *r* with no contacts, 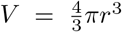 and *A*_exposed_ = 4*πr*^2^, giving *ϕ* = *r/*3. *ϕ* is normalized to the isolated-cell reference geometry (*ϕ* = 1, Fig. 1(E), schematic (i)). Cell-cell contacts reduce *A*_exposed_ while *V* is conserved, raising *ϕ* above 1 (Fig. 1(E), schematic (ii)).

#### Intracellular Feedback Loop

Intracellular feedback is incorporated by modeling the endocytosis rate *β* as a sigmoidal function of the internalized receptor pool:

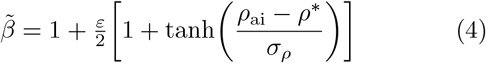

where *ρ*\* is the feedback threshold, *σ*_*ρ*_ the steepness, and *ε* the feedback strength. When *ρ*_ai_ *< ρ*\*, feedback is negligible (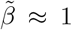, basal endocytosis) but when *ρ*_ai_ *> ρ*\*, feedback becomes strong (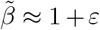, enhanced endocytosis), creating a non-monotonic *ρ*_a_(*c*) that peaks at *c*\* and falls at higher *c* (Fig. 1(E)), as derived in the next subsection. Note that the exact functional form of the feedback 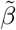 given in Eq. (4) is not important, and the results are similar as long as it has a threshold-like (highly nonlinear) dependence on the density of internalized receptors *ρ*_*ai*_.

#### Geometric Parameter and *c*\* Scaling

At steady state, Eqs. (1–4) yield:

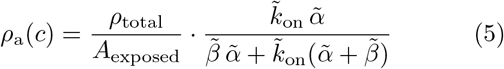

with 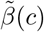 determined self-consistently through Eq. (4) via 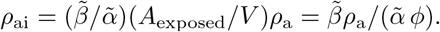.

The chemotactic sensitivity, *χ*(*c*) = *dρ*_a_*/dc*, determines the migration direction with chemoattraction when *χ >* 0 (migration up the gradient) and chemorepulsion when *χ <* 0 (migration down the gradient). In our model the local density of active receptors on the cell membrane determines the local strength of the protrusive activity of the cytoskeleton. The difference between this activity across the cell due to the chemokine concentration gradient determines the direction of the cell migration. Without the internalization feedback (*ε* = 0), *ρ*_a_(*c*) increases monotonically and cells undergo positive chemotaxis. With feedback, *ρ*_a_(*c*) is non-monotonic (Fig. 1(E)). At low *c*, feedback is weak and ligand binding dominates (*χ >* 0), while, above a critical concentration *c*\* (where *dρ*_a_*/dc* = 0), feedback-accelerated endocytosis outpaces binding, depleting surface receptors and reversing polarity (*χ <* 0, chemorepulsion). This polarity inversion (a sign change in *χ*(*c*)) is the mechanistic origin of single-cell chemorepulsion in this model.

### (ii) Agent-Based Model for Cell Clusters

#### ABM with coarse-grained cell switching

The single-cell receptor model predicts a sign change in cell migration above *c*\*, but does not describe how this effect propagates to multicellular dynamics. We therefore introduce an agent based model (ABM) to bridge this gap (Fig. 2). The model consists of *N* cells (soft disks in 2D) that interact mechanically and switch between chemoattractive and chemorepulsive polarity states. The forces that act on a cell are active self propulsion (due to motility), steric exclusion and adhesion to neighboring cells with its orientation being influenced by those of its neighbors and the locally sensed gradient direction. A cell’s motion is governed by overdamped Langevin dynamics. All model details are described in the Materials and Methods section (and Fig. M1a).

**FIG. 2.**
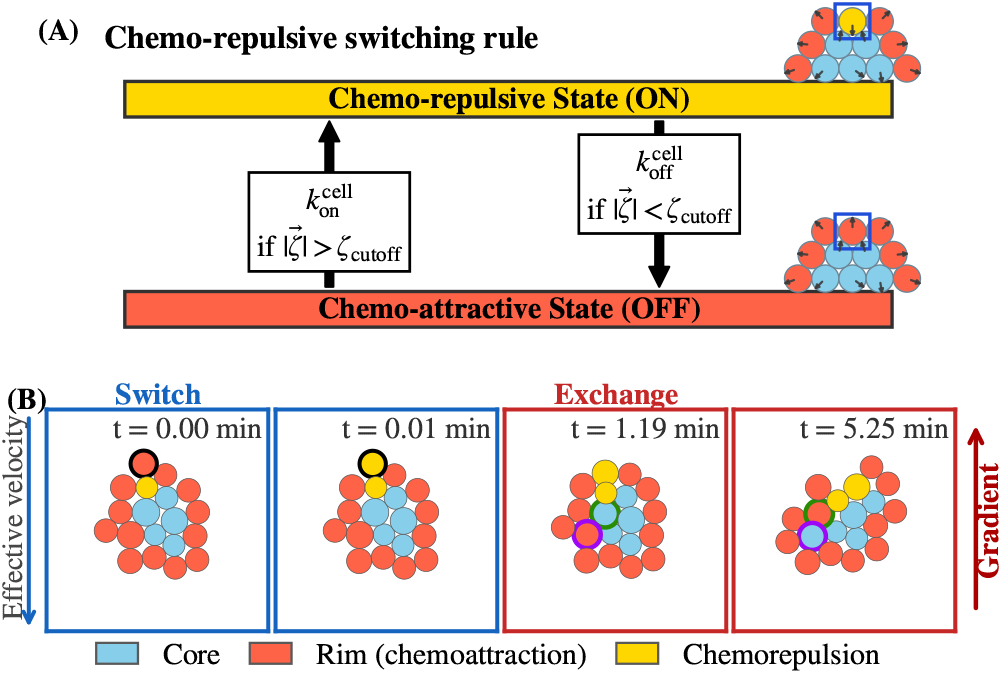
Chemorepulsive switching rule and simulation time series. **(A) Switching rule**. Cells switch between chemoattractive (OFF, orange–red) and chemorepulsive (ON, yellow) states via Heaviside-gated rates [Eq.(10)]. Core cells (blue, 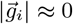) remain attractive; leading-edge rim cells exceeding ζ_cutoff_ switch to repulsive (yellow). **(B) Time series (***N* ≈ 19, *t* = 0.00, 0.01, 1.19, 5.25 **min)**. *Switch* (*t* = 0 → 0.01 min): the highest-exposed rim cell (highlighted in black) crosses ζ_cutoff_ and switches to the repulsive state before any cluster displacement, confirming a receptor-level trigger. *Exchange* (*t* = 1.19 → 5.25 min): as the cluster deforms, cells exchange between rim and core (highlighted purple) or core to rim (highlighted in green). Newly exposed cells may switch; formerly repulsive rim cells shielded by neighbors revert to the attractive state, preventing permanent accumulation of repulsive cells at the leading edge.

The ABM does not resolve receptor populations explicitly at the single cell level. Instead, each cell *i* carries a coarse-grained polarity variable *s*_*i*_ ∈ {+1, − 1}, which is either chemoattractive (*s*_*i*_ = +1) or chemorepulsive (*s*_*i*_ = − 1) (Fig. 2(A)), encoding only the sign of the analytically predicted polarity inversion. A change in the sign of *s*_*i*_ affects the cell’s motion in the ABM by flipping the sign of the locally sensed gradient direction which in turn influences the cell’s preferred orientation.

#### Local sensing variable and switching criterion

We define a local sensing variable, 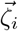, combining each cell’s geometric access to the chemokine gradient 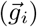 with saturating receptor activation (*χ*) and overall sensing strength (*ψ*)

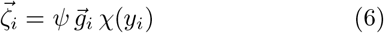

The local open-area vector, 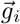, captures the direction and magnitude of the most exposed area of the cell boundary (see Materials and Methods section (Fig. M1b)). A core cell uniformly surrounded by neighbors has 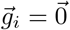 (no net open area), while a rim cell with a large exposed arc has a large 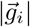 (details in the Materials and Methods section). This models cell–cell contacts blocking receptor binding in the contact zone, with larger exposed arcs allowing greater receptor engagement and directional sensing.

The receptor saturation factor *χ*(*y*_*i*_) is the bound fraction of CCR7 receptors at position *y*_*i*_, modeled by Langmuir adsorption:

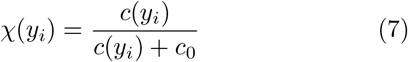

where *c*_0_ is the half-saturation concentration. The local CCL19 concentration is

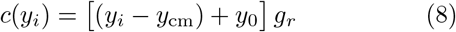

where *y*_cm_ is the cluster center of mass (CM) *y*-position, *g*_*r*_ is the slope of the spatial chemical gradient, and *y*_0_ is the CM offset from the gradient origin (*y*_0_ *>* | *y*_*i*_ − *y*_cm_ | ensures non-negative concentrations). Here, *c*_CM_ = *y*_0_ *g*_*r*_ is the average CCL19 concentration at the cluster CM.

In the coarse-grained ABM, a cell transitions to the chemorepulsive state when its local sensing variable exceeds a universal threshold (see Fig. 2(A)):

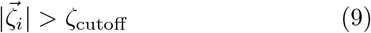

This cutoff arises from the analytical receptor model which predicts that the sensing variable evaluated at the polarity-inversion point, ζ(*c*\*(*ϕ*)), is approximately constant across all volume-to-exposed area ratio (*ϕ*) values, a direct prediction of the analytical model. This near-constancy emerges from two opposing *ϕ* dependencies: as *ϕ* increases, the open area 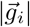 decreases (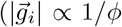, since more neighbors occlude more membrane), while simultaneously the polarity-inversion concentration *c*\* increases (*c*\* ∝ *ϕ*, SI Appendix, Section S1). At the inversion point these two effects cancel: the product 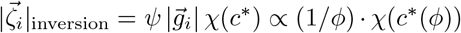 is nearly constant because the decreasing geometry factor (1*/ϕ*) is offset by the increasing receptor saturation *χ*(*c*\*(*ϕ*)) (SI Appendix, Fig. S1D; Section S1). A single ζ_cutoff_ can therefore be used for all cells in all cluster sizes without refitting (calibration and isolated-cell details are provided in SI Appendix, Section S1).

Transitions between polarity states are stochastic (see Fig. 2(A)), governed by Heaviside-gated switching rates:

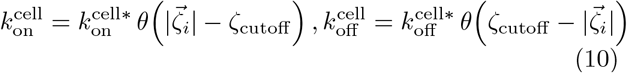

where 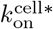 and 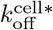 are the maximal switching rates and *θ* is the Heaviside step function. Above ζ_cutoff_, a cell switches to the repulsive state (*s*_*i*_ → − 1) with rate 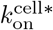; below ζ_cutoff_ (or when 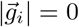, i.e., a core cell), it returns to the attractive state (*s*_*i*_ → +1) with rate 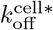. This stochastic formulation reflects unresolved biochemical noise, cell-to-cell heterogeneity, and receptor trafficking fluctuations. In this model, polarity inversion is probabilistic at the individual cell level, and its collective consequences emerge through mechanical interactions, spatial rearrangements, and boundary-bulk turnover within the cluster (Fig. 2(B)).

#### ABM switching rate calibration

CCR7 internalization and recycling occur on minute-to-hour timescales [35, 39, 41, 42]. We calibrate the ABM switching rates directly to these CCR7 trafficking kinetics. 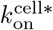 is set by the CCR7 endocytosis timescale (*τ*_intern_ ≈ 20–30 *min*):

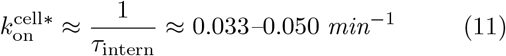

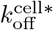 is set by the CCR7 recycling timescale (*τ*_recycle_ ≈ 1 *h*):

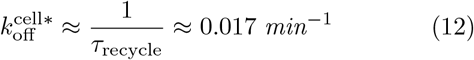

The full ABM implementation details incorporating this switching dynamics as well as the equations of motion of the particles, integration scheme, and boundary conditions are provided in the Materials and Methods section.

## RESULTS

We begin with the experimental observation that motivates this study. Time-lapse microscopy of JVM3 (CLL) cells in a 0–500 ng/mL CCL19 gradient reveals a behavioral paradox: single cells migrate *away* from the chemokine source (chemorepulsion, FMI = − 0.65, *n* = 43 tracks), whereas clusters of *N* ≥ 20 cells migrate *toward* it (chemoattraction, FMI = +0.49, *n* = 23 tracks, Fig. 1A–B). Both populations experience the same extracellular gradient and express the same CCR7 receptor, indicating that the sign reversal cannot arise from receptor identity alone.

### Receptor Trafficking Feedback Drives Single-Cell Chemorepulsion

The analytical receptor trafficking model (Fig. 1D) predicts a non-monotonic surface receptor density *ρ*_*a*_(*c*) with a switching threshold at a critical concentration *c*\* (Fig. 1E). Below *c*\*, cells remain chemoattractive (Fig. 1E, shaded blue), whereas above *c*\* the migration polarity reverses(Fig. 1E, shaded red). This reversal emerges because intracellular feedback-driven endocytosis progressively depletes surface receptors at high ligand concentrations.

Two independent experimental observations validate this molecular switching mechanism for isolated JVM3 cells. First, inhibiting dynamin-dependent endocytosis converts single-cell chemorepulsion into chemoattraction [31], confirming that active endocytosis is required for the repulsive state. Second, concentration-sweep experiments show that, with the gradient slope fixed, increasing the mean CCL19 concentration progressively shifts isolated JVM3 cells from chemoattraction to chemorepulsion (see red bars in Fig. 3E). Specifically, moving the CCL19 range from 0–100 ng/mL (average ≈ 50 ng/mL, below *c*\*) to 400–500 ng/mL and then 1400– 1500 ng/mL drives the transition predicted by the non-monotonic *ρ*_*a*_(*c*) curve in Fig. 1E. Importantly, cells exposed to a spatially uniform high-concentration CCL19 field exhibit random motion rather than directed repulsion(Figure 1B–C, [31]), demonstrating that receptor saturation alone is insufficient to explain the switch. Instead, polarity reversal requires a gradient-induced asymmetry operating above *c*\*. Together, these observations establish the molecular basis of single-cell chemorepulsion and identify the switching threshold *c*\* that cluster geometry modulates at the collective level.

**FIG. 3.**
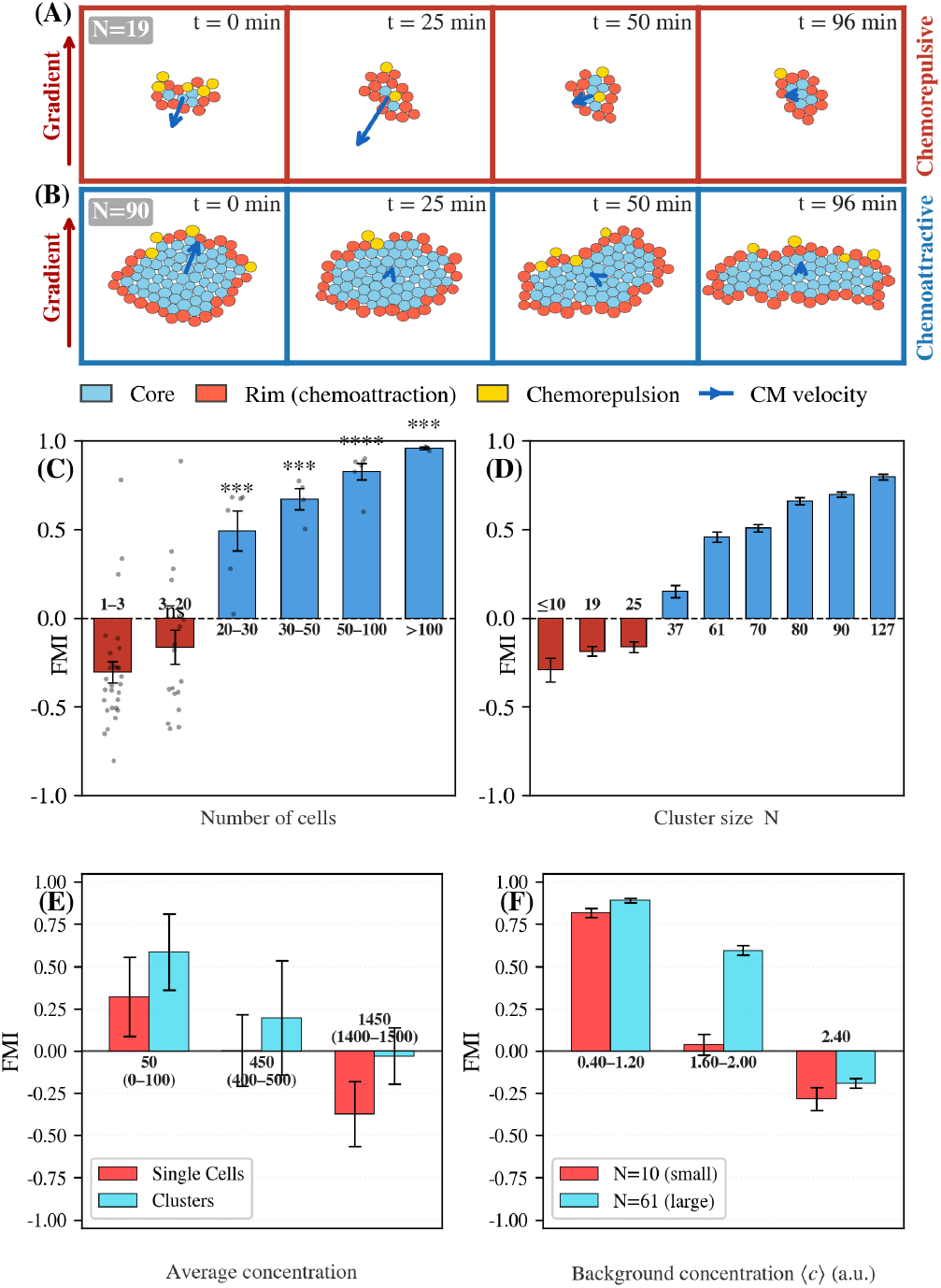
Size-dependent chemotactic switch: simulation and experimental validation. **(A)** Simulation snapshots of *N* = 19 (chemorepulsive) at *t* = 0, 25, 50, 96 min. Downward CM velocity arrows confirm persistent migration away from the source. **(B)** Simulation snapshots of *N* = 90 (chemoattractive) at the same time points. The cluster elongates and migrates toward the source (upward arrows), sustained by rim-core fluidity. **(C)** Experimental FMI vs. cluster size (CCL19 0–500 ng/mL, JVM3): ~ 1–3 cells (*n* = 29), ~ 3– 20 (*n* = 18), ~ 20–30 (*n* = 6), ~ 30–50 (*n* = 4), ~ 50–100 (*n* = 6), *>*100 (*n* = 3). FMI becomes significantly positive at ~ 20–30 cells (***) and increases monotonically, mirroring panel (D)). **(D)** Model FMI vs. *N* (*N* = 7–127; *n* = 18– 40 runs). Clusters with *N* ≲ 25 show negative FMI (red); *N* ≤ 37 show positive FMI rising to ≈ +0.8 (blue). Error bars: SEM. **(E)** Experimental FMI vs. average CCL19 slab concentration (fixed gradient slope; each tick shows the mean concentration with its slab range in parentheses; JVM3). Single cells (red) switch to chemorepulsion as average slab concentration increases; cluster FMI (blue) also declines with concentration but remains near zero at the highest slab tested, consistent with the model prediction in panel (F) that elevated ⟨c⟩ more strongly affects small than large clusters. **(F)** Model FMI vs. average ligand concentration at the cluster CM (⟨c⟩= *y*_0_ *g*_*r*_; see Methods) for small (*N* = 7, 10; red) and large (*N* = 37, 61; blue) clusters. Small clusters cross the FMI sign boundary at elevated ⟨c⟩; large clusters remain chemoattractive.

### Cluster Geometry Suppresses Chemorepulsion by Raising the Switching Threshold

We next asked how cell–cell contacts alter this molecular threshold. Cell–cell contacts raise the volume-to-exposed-area ratio *ϕ* (= *V/A*_exp_) by reducing exposed membrane while conserving volume. As *ϕ* increases, *c*\* shifts monotonically to higher concentrations (Fig. 1E, SI Appendix, Fig. S1A,C). At *ϕ* = 1.5 (roughly one third of membrane occluded), the model predicts *c*\* = 306. At *ϕ* = 2.0 (half the membrane occluded), *c*\* = 501, and at *ϕ* = 5.0, shifting the polarity-inversion point beyond the gradient range in the model.

The *c*\* − *ϕ* relationship is approximately linear (SI Appendix, Fig. S1C).This predicts that ζ(*c*\*(*ϕ*)) is approximately constant across physiological *ϕ* values (SI Appendix, Fig. S1D), allowing a single ζ_cutoff_, calibrated from isolated-cell data, to apply across cluster sizes without re-fitting. Robustness to this choice is confirmed by a ζ_cutoff_ sweep (SI Appendix, Fig. S4).

Furthermore, as *ϕ* increases, the chemorepulsive concentration window, defined by *dρ*_*a*_*/dc <* 0, narrows (SI Appendix, Fig. S1B), consistent with the weaker and less persistent repulsion observed at intermediate cluster sizes experimentally [31]. Thus, cluster geometry converts receptor trafficking into a local switching rule that can be embedded in the ABM through each cell’s instantaneous exposed-area geometry.

### The ABM Reproduces the Chemorepulsion–Chemotaxis Switch and Reveals the Collective Mechanism

The ABM reproduces the key experimental observation: large clusters (*N* ≥ 20) migrate toward the CCL19 source while small clusters migrate away (Fig. 1A–B), consistent with the sign reversal reported by Malet-Engra et al. [31]. We first examine the early-time dynamics, which reveal that polarity switching is receptor-driven. Simulation snapshots of a small cluster (*N* ≈ 19 cells) display a two-phase dynamics (Fig. 2B). Between *t* = 0.00 and *t* = 0.01 min, the leading-edge rim cell exposed to the highest local CCL19 concentration crosses ζ_cutoff_ and transitions from the chemoattractive (orangered) to the chemorepulsive (yellow) state. Over longer timescales (*t* = 1.19–5.25 min), the cluster deforms and cells exchange dynamically between the rim and the core (Fig. 2B). Chemorepulsive cells at the periphery tend to push themselves towards the center of the cluster. Cells displaced outward inherit the high-signal rim environment and may switch into the repulsive state, whereas previously repulsive rim cells shielded by neighboring cells revert to the chemoattractive state. These dynamics redistribute the repulsive cells rather than allowing them to accumulate permanently at the leading edge.

### FMI responds to cluster size, gradient steepness, and ambient concentration

We now show that cluster size determines the sign of collective migration at fixed gradient steepness. Contrasting large (*N* ≈ 90) and small (*N* ≈ 19) cluster migration velocities (shown in (Fig. 3A–B)) directly reveals this size dependence. The large cluster elongates perpendicular to the gradient and migrates persistently toward the CCL19 source (positive CM velocity), whereas the small cluster migrates away from the source (negative CM velocity).These migration directions emerge from receptor-level switching coupled to collective rim-core redistribution, producing the size-dependent FMI sign reversal observed experimentally. To quantify this size dependence, we compute FMI as a function of cluster size across the full range *N* = 7–127. For a single calibrated parameter set (ζ_cutoff_, *g*_*r*_ = 0.008), the ABM FMI reproduces the experimental measurements of Malet-Engra et al. [31] across all tested cluster sizes (Fig. 3C–D). Small clusters (*N* ≲ 25) exhibit FMI *<* 0 (chemorepulsion, red bars), whereas larger clusters exhibit FMI *>* 0 (chemoattraction, blue bars), with FMI increasing toward ≈ +0.8 at large *N*. The simulated FMI sign reversal between *N* ≈ 25 and 37 cells is consistent with the experimentally observed transition near ~ 20–30 cells [31]. The same parameter set also reproduces independent observables across cluster sizes, including center-of-mass migration speed (≈ 4 *µ*m/min, SI Appendix, Fig. S3), rim-to-core exchange time, and leader-cell dwell time (SI Appendix, Fig. S2A,B). Both exchange time and dwell time decrease with increasing *N*, as the growing rim population accelerates turnover, while remaining consistent with the experimentally reported 8–15 min leader dwell time [31].

Leader-cell dwell time further provides a dynamical interpretation of experimentally observed CCR7 surface fluctuations in leader cells [31]. During prolonged exposure at the rim, surface CCR7 is progressively depleted through endocytosis, whereas replacement by interior cells restores receptor levels. Each fluctuation cycle therefore corresponds to one rim-core exchange event (full analysis in SI Appendix, Section S2).

The calibrated model predicts a second axis of control: average background CCL19 concentration independently drives the migration switch even at fixed cluster size. Increasing ambient CCL19 drives small clusters (*N* = 7– 10) across the FMI sign boundary, whereas larger clusters (*N* = 37–61) remain predominantly chemoattractive (Fig. 3F). This prediction is confirmed experimentally for clusters (Fig. 3E, blue), where FMI progressively decreases as the ambient CCL19 concentration increases at fixed gradient, and is barely negative even at 1400– 1500 ng/mL.

The corresponding single-cell measurements from the same experiment show isolated cells switching from chemoattraction to chemorepulsion as ambient concentration increases (Fig. 3E, red). Together, the two curves in Fig. 3E show that increasing ambient CCL19 drives isolated cells above their switching threshold, whereas cluster cells remain protected, staying below threshold because of the *ϕ*-dependent shift of *c*\*.

### The Chemorepulsive Fraction Predicts Collective Migration Direction

Having established that collective FMI responds to cluster size, gradient steepness, and ambient concentration, we next asked whether a single internal state variable predicts these dependencies. Sweeping across both cluster size *N* and gradient steepness in the ABM reveals a two-dimensional phase diagram (Fig. 4A). The chemorepulsive regime (FMI *<* 0, red) occupies the small-*N*, steep-gradient corner, whereas the chemoattractive regime (FMI *>* 0, blue) occupies large *N* and/or shallow gradients. The FMI = 0 (black line on (Fig. 4A)) boundary shifts to larger *N* as gradient steepness increases, indicating that stronger gradients recruit more rim cells past ζ_cutoff_ and therefore require larger clusters to maintain net chemoattraction. This model prediction is consistent with the gradient-steepness dependence reported in Malet-Engra et al. [31].

**FIG. 4.**
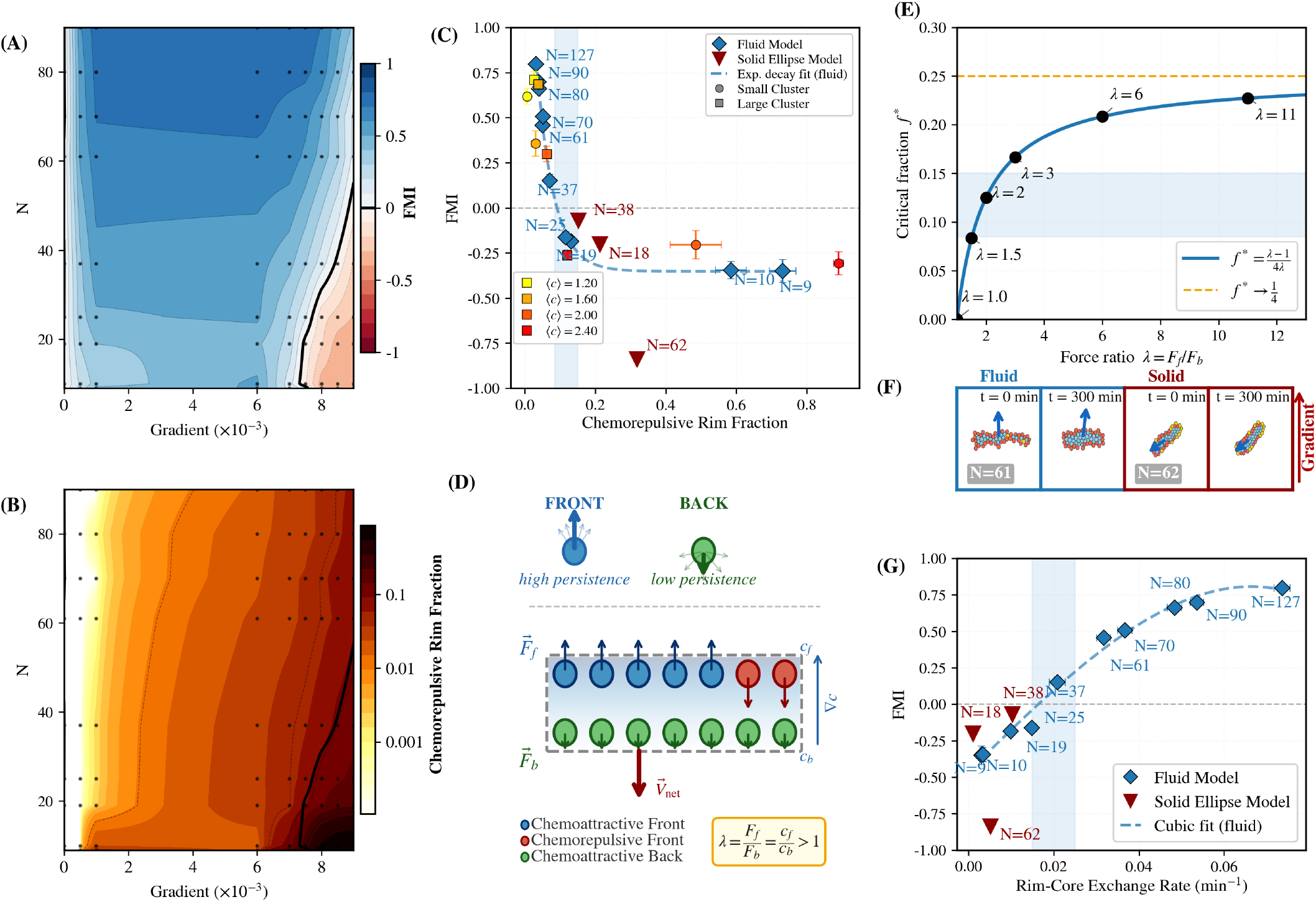
Phase diagram, scaling, and critical threshold for the chemorepulsion–chemotaxis switch. **(A)** FMI heatmap (*N* = 9–90, ∇*c* = 0–9 *×* 10^−3^). Chemorepulsive regime (red) occupies the small-*N*, steep-gradient corner, whereas the chemotactic regime (blue) occupies larger *N* and/or shallower gradients. The FMI= 0 boundary shifts to larger *N* with increasing gradient steepness. **(B)** Chemorepulsive fraction *f* heatmap over the same (*N*, ∇*c*) space. High chemorepulsive fraction mirrors the repulsive FMI corner; together, (A) and (B) show that collective migration direction is governed by the accumulation of chemorepulsive cells at the cluster rim. **(C)** FMI versus chemorepulsive rim fraction *f*. Fluid clusters (blue diamonds) maintained below the critical fraction *f* * (shaded) exhibit positive FMI, whereas the solid ellipse model accumulates larger chemorepulsive fractions and remains strongly chemorepulsive. The dashed blue line shows an exponential decay fit, *A e*^−*λf*^ + *C*, through the fluid-model points. Concentration-sweep points (⟨c⟩ = 1.20–2.40 a.u., *g*_*r*_ = 0.008) are coloured by average concentration ⟨c⟩ = *y*_0_ *g*_*r*_ (warm colorscale; see legend) for small clusters (*N* = 7, 10 averaged; circles) and large clusters (*N* = 37, 61 averaged; squares), showing that increasing *c* drives both cluster classes toward higher chemorepulsive fractions and lower FMI, consistent with the gradient-sweep trend. **(D)** The upper persistence sketch shows that front cells have higher persistence, whereas back cells have lower persistence, providing a mechanistic basis for stronger effective front forces. Minimal toy-model schematic for the critical switching threshold. Front cells experience higher signal than back cells, giving a force asymmetry *λ* = *F*_*f*_ */F*_*b*_ = *c*_*f*_ */c*_*b*_ *>* 1. In the lower two-row schematic, the front row can therefore contain both chemoattractive and chemorepulsive cells, while the back row remains chemoattractive. The net migration velocity *V*_net_ reflects the competition between front- and back-derived forces. **(E)** Analytical prediction for the critical chemorepulsive fraction as a function of force asymmetry, *f* * = (*λ* − 1)*/*(4*λ*). The curve increases with *λ* but remains bounded by 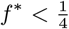, implying that a minority of front chemorepulsive cells is sufficient to reverse collective migration. The simulated value *f* * ≈ 0.10 lies well within this bound. **(F)** Fluid model (*N* = 61) at *t* = 0 and 300 min: the cluster migrates toward the CCL19 source through ongoing rim-core exchange. Solid ellipse (*N* = 62, 4-layer: 15–16–16–15), rate-matched but exchange-suppressed: chemorepulsive cells accumulate at the leading edge and the cluster migrates away from the source. **(G)** FMI versus rim-core exchange rate. Fluid-model clusters (blue diamonds, *N* = 9–127; *y*_0_ = 250, *g*_*r*_ = 0.008) show a sharp transition from negative to positive FMI as exchange increases, whereas the solid ellipse model (red triangles) remains at low exchange and negative FMI. The dashed blue line shows a cubic polynomial fit through the fluid-model points.

The fraction of all cells in the cluster that are chemorepulsive, *f*, mirrors this pattern (Fig. 4B). High *f* concentrates in the same small-*N*, steep-gradient corner, indicating that collective migration direction is set by the balance between repulsive rim cells and shielded chemoattractive cells. Across four gradient steepnesses, *f* decays with cluster size as ~ *N* ^−0.6^–*N* ^−1.2^ (SI Appendix, Fig. S5A), with steeper gradients exceeding the *N* ^−0.8^ geometric perimeter scaling. The steeper decay at higher gradient steepness reflects that more rim cells cross the switching threshold at any given *N*, increasing *f* at each cluster size and making the decline with *N* more abrupt. However, the underlying reason *f* decreases with *N* in all cases is geometric. Larger clusters have proportionally fewer rim cells relative to their total size, regardless of gradient steepness.

Plotting FMI directly against the instantaneous chemorepulsive fraction for *N* = 19–61 across varying gradient steepnesses, with concentration-sweep data overlaid, reveals a single monotonically decreasing curve (Fig. 4C). Thus, *f* collapses the effects of cluster size, gradient steepness, and ambient ligand concentration onto one scalar predictor of collective migration direction. From simulations, the sign change occurs at a critical fraction *f* * ≈ 0.10: Clusters with *f < f* * are chemoattractive (FMI *>* 0), whereas clusters with *f > f* * are chemorepulsive (FMI *<* 0). This computationally derived threshold is independently confirmed by an analytical force-balance model, described next (Eq. 13).

A minimal two-layer force-balance model provides an independent bound on the repulsive fraction required to reverse collective migration (Fig. 4D–E, SI Appendix, Section S5). The model consists of two rows of *n* cells each (Fig. 4D). Front cells experience a higher local CCL19 concentration than back cells. Higher concentration drives higher persistence in front cells, so on average each front cell generates a stronger propulsive force *F*_*f*_. Back cells experience lower concentration, which gives them lower persistence and therefore a weaker propulsive force *F*_*b*_ per cell. Together these give the force asymmetry *λ* = *F*_*f*_ */F*_*b*_ = *c*_*f*_ */c*_*b*_ *>* 1 (Fig. 4D, upper sketch). The back row remains chemoattractive throughout. The front row can contain both chemoattractive and chemorepulsive cells depending on whether local concentration exceeds *c*\*. The net cluster velocity 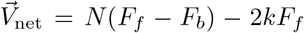, where *k* is the number of repulsive cells in the front row, reflects the competition between the forward drive from chemoattractive cells and the backward pull from chemorepulsive front cells (Fig. 4D, lower sketch). Writing *f* = *k/*(2*N*) as the fraction of all cells that are repulsive and setting *V*_net_ = 0 gives the critical chemorepulsive fraction

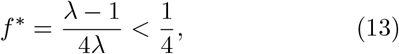

The curve in Fig. 4E shows that *f* * rises monotonically with *λ*. Stronger gradient asymmetry means the remaining attractive front cells push harder forward, so a larger repulsive fraction is needed to tip the balance. Nevertheless, 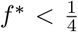 for all *λ >* 1, confirming that collective reversal always requires fewer than one quarter of all cells to switch. The computationally derived *f* * ≈ 0.10 falls well within this bound. The small magnitude of *f* * reflects the force asymmetry. Because front cells generate a stronger force *F*_*f*_ *> F*_*b*_, switching even a small number of front cells from chemoattractive to chemorepulsive shifts the force balance disproportionately, since each switched cell converts a +*F*_*f*_ contribution into − *F*_*f*_, a net change of 2*F*_*f*_ in the backward direction.

### Cluster Fluidity Enables Collective Chemoattraction by Keeping *f* Below the Critical Fraction

Rim-core exchange is the collective mechanism that complements geometric shielding: by continuously cycling cells between the cluster rim and interior, large fluid clusters maintain *f* below *f* * ≈ 0.10 (Eq. 13). We finally asked why large clusters maintain *f < f* * while small clusters do not. To isolate the role of cluster fluidity, we compared the current full fluid model against a solid ellipse control in which internal positional exchange is suppressed while the turn-on rate 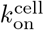 is matched (Fig. 4F; SI Appendix, Section S6). Cluster fluidity in the model arises from CIL-driven motility of interior cells, which produces stochastic rim-core displacements.

Fluidity changes migration direction only once clusters are large enough for rim-core exchange to dilute *f* below *f* *. For small clusters (*N* ≈ 19), both the fluid and solid models are chemorepulsive (FMI = − 0.216 and 0.200, respectively; Fig. 4G). For intermediate clusters (*N* ≈ 37), the fluid model crosses weakly into chemoattraction (FMI = +0.060), whereas the solid model remains chemorepulsive (FMI = − 0.069). For large clusters (*N* ≈ 61), the contrast is dramatic: the fluid model achieves FMI = +0.309, while the solid model collapses to FMI = − 0.836. Without exchange, every additional cell exposed to the steep gradient permanently occupies the repulsive front, so larger solid clusters accumulate more repulsive force with no compensating redistribution. Thus, fluidity becomes decisive only beyond the size at which exchange can reduce *f* below the critical fraction. In small clusters, the rim constitutes a proportionally large fraction of total cells, so even rapid exchange quickly returns cells to the rim environment where they re-enter the repulsive zone. The dilution effect cannot outpace re-exposure until the cluster is large enough that the core acts as a sufficient reservoir.

Plotting FMI directly against rim-core exchange rate reveals a critical fluidity threshold at ≈ 0.02 exchanges per unit time (Fig. 4G), corresponding to the exchange rate at which *f* crosses *f* * ≈ 0.10 (Fig. 4C). The exchange rate is the mechanistic control variable and *f* is the resulting state variable. Below this value, clusters are chemorepulsive. Above it, FMI increases steeply with both exchange rate and *N*. Fluid-model clusters lie to the right of this threshold. Solid ellipse controls remain near zero exchange rate with strongly negative FMI. Thus, rim-core exchange provides the dynamical control that keeps *f* below the critical fraction in large clusters, whereas exchange-suppressed clusters accumulate repulsive rim cells and migrate away from the source. Together with the *ϕ*-dependent shift of *c*\*, this identifies geometry and fluidity as the two collective mechanisms that convert single-cell chemorepulsion into cluster-level chemoattraction.

## DISCUSSION

Our study identifies three nested mechanisms at the molecular, cellular, and collective levels that jointly resolve the paradox of CLL single cells chemorepulsing in large gradients of chemoattractant, while clusters of the same cells chemoattract in the same gradients. At the molecular level, intracellular positive feedback on receptor endocytosis creates a polarity-inversion concentration threshold above which activated surface receptors are depleted faster than ligand binding can replenish them. At the cellular level, cell-cell contacts raise the volume-to-exposed-area ratio (*ϕ*), diluting the internalized receptor pool and shifting the critical chemorepulsion concentration beyond the physiological gradient window. At the collective level, rim-core cell exchange keeps the chemorepulsive cell fraction at the cell rim below the critical fraction, sustaining net chemoattraction in large clusters.

We show that no new molecular mechanisms are required to explain the observations, with intracellular dynamics remaining identical in single cells and cluster cells. At the single cell level, our model shares some similarity with a recent model [38] where the concentration-dependent chemorepulsion switch in isolated cells is generated by a proposed downstream activator-inhibitor circuit as opposed to the direct feedback from activated receptor internalization in our model. While the same qualitative behavior is observed for single cells, this model does not address cluster chemotaxis. In our model, the volume-to-exposed-area changes as cell-cell contacts occlude membrane surface, reducing receptor internalization relative to cell volume and suppressing the positive-feedback loop. This geometric mechanism operates entirely through the physical consequence of cell-cell contact. It should be noted that cell-cell contacts can additionally trigger junction-associated signaling, for example through cadherin-mediated pathways [49] or Notch [50], which in turn can affect cytoskeletal dynamics and chemotactic response. Our framework does not exclude such contributions. Single-cell chemorepulsion is a general regulatory feature seen in other contexts such as neutrophil reverse migration away from inflamed tissue [7]. This suggests that feedback-driven repulsion is a conserved regulatory mode across immune cell types. Clustering suppresses this escape mechanism, allowing the collective to maintain forward migration. We therefore propose that receptor endocytosis functions not merely as a degradation pathway but as a tunable directional switch whose threshold is collectively controlled by cluster geometry.

The collective mechanism adds a second, dynamic layer of control. Rim-core fluidity continuously redistributes chemorepulsive rim cells away from the leading edge even under steep gradients. We isolated this effect by comparing fluid and solid clusters with similar rates of cellular chemorepulsive switching. Clusters without internal exchange lock into chemorepulsion at large sizes, while fluid clusters of identical size maintain positive FMI (Fig. 4F–G). This demonstrates that rim-core fluidity is a necessary and sufficient physical property for collective chemoattraction at large cluster sizes.

In many contexts, receptor internalization can produce a chemokine gradient as established for CCR7 in dendritic cells [35], where CCL19 internalization reshapes the extracellular gradient to enable self-generated guidance. Both effects stem from the same process, endocytosis, but operate at different spatial scales. We test this directly from the existing data. Cells collectively depleting CCL19 would produce more persistent and elongated tracks in any direction, even without an imposed gradient [36, 37]. We find, however, that track shapes and persistence are indistinguishable between the uniform-field and no-CCL19 conditions in Malet-Engra et al. [31], whereas gradient conditions produce markedly more directional tracks. We conclude that self-generated gradient guidance is not significant in this open microfluidic geometry and that the behavioral switch is accounted for by cell-cell occlusion (*ϕ*) and cluster fluidity.

Our three-level mechanism carries direct implications for CLL lymph node homing. CCL19 is constitutively expressed in lymph nodes, the primary site of CLL proliferation [4]. Steeper gradients and higher average concentration both push more rim cells above the threshold for chemorepulsion (ζ_cutoff_), shifting the chemorepulsion-to-chemotaxis boundary to larger cluster sizes (Fig. 4A) and progressively eroding cluster FMI (Fig. 3E–F). This may explain why CLL cells accumulate preferentially as clusters in lymphoid tissue, potentially increasing the likelihood of metastasis by circulating tumor clusters.

In summary, receptor trafficking feedback, cluster geometry, and collective rim-core fluidity form three nested mechanisms that quantitatively account for how cluster geometry gates chemotactic decision-making in cells. These findings extend to many other biological systems where cells migrate as individuals and as clusters along chemical gradients,including immune cell responses, embryonic morphogenesis, and cancer progression.

## MATERIALS AND METHODS

### Cell Line and Reagents

The human CLL-derived cell line JVM3 was obtained from the Deutsche Sammlung von Mikroorganismen und Zellkulturen (DSMZ) and authenticated by B cell surface marker analysis, CCR7 expression, and mycoplasma testing. Recombinant human CCL19 was purchased from PeproTech; Alexa Fluor 647–labeled CCL19 was purchased from Almac. Both were aliquoted and stored according to the manufacturers’ instructions.

### Chemotaxis Assay and Imaging

Cell migration in CCL19 gradients was assessed by video microscopy using collagen IV–coated 2D chemotaxis *µ*-slides (ibidi), as previously described [31]. Cells (0.5 *×* 10^6^ in 10 ml culture medium) were loaded into the central channel and allowed to adhere at 37^*◦*^C for 30 min. Chemokine gradients were established according to the manufacturer’s protocol and verified for linearity using 10% dextran–fluorescein isothiocyanate solution. Cell migration was recorded under environmental control (temperature and CO_2_) at one frame per 2 min using MetaMorph software (Molecular Devices) [31] on either a Zeiss Axiovert 200 microscope (2.5 *×* objective, coolSNAP HQ CCD camera, Photometrics) or a Nikon Eclipse TE2000-E microscope (4 *×* /0.13 NA objective, Cascade II 512 CCD camera, Photometrics).

### Image Analysis

Cluster shape and orientation were quantified from time-lapse images using the built-in Measure function in ImageJ. Each cluster was fitted to an ellipse and the orientation angle of the major axis was extracted. Cluster breaking events were identified by monitoring cluster size over time.

## AGENT-BASED MODEL: FULL SIMULATION FRAMEWORK

The agent-based model (ABM) represents each CLL cell as a self-propelled disk of radius *r*_*i*_ moving in a two-dimensional gradient of CCL19. Each cluster of *N* cells is assigned a distribution of radii with a polydispersity of ~10%, reflecting the natural size variability of CLL cells and suppressing artificial crystalline ordering. The mechanical core of the model (CIL, Lennard-Jones interaction, cohesive spring, velocity alignment, and stochastic noise) was introduced by Copenhagen et al. [22], whose parameters were calibrated by matching the three canonical gradient-free migration modes (rotating, running, and random) to experimental observations of CLL clusters [31]. Gradient sensing 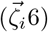, *χ*(Eq 7), *c*(*y*)(Eq 8)) was subsequently incorporated by Sanoria et al. [24], with sensing parameters calibrated against gradient-driven cluster shape instabilities and migration efficiency measurements [31]. The present model adds chemorepulsion switching (*s*_*i*_, ζ_cutoff_) as the new ingredient. The dynamics are overdamped: inertia is neglected and the instantaneous velocity equals the sum of all forces acting on cell *i* (Fig M1(a)):.

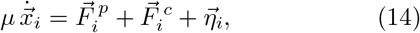

where *µ* is a damping coefficient, 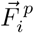 is the self-propulsion (motility) force, 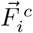 is the cell-cell contact force (Lennard-Jones interaction plus cohesive spring), and 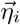 is a stochastic noise term. We integrate (14) with a forward-Euler step of size Δ*t* = 0.01 sim. units (= 0.005 min). The subsections below define each ingredient in sequence, starting with self-propulsion (CIL speed law and polarity direction), followed by the geometric sensing quantities 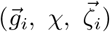, then mechanical cell-cell forces (LJ and cohesive spring), stochastic noise, and finally the polarity switching rule that couples 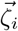 to *s*_*i*_ and thereby to the direction of migration.

### Self-Propulsion Force

The self-propulsion speed *p*_*i*_ of each cell is governed by contact inhibition of locomotion (CIL) [22]: cell propulsion is smaller when cells are confined by neighbors, with rim cells moving faster than core cells because they have more open space available to form protrusions. The propulsion speed is taken to be:

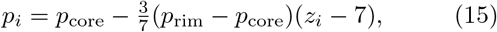

where *p*_rim_ and *p*_core_ are the propulsion speeds of rim and core cells respectively, and *z*_*i*_ is the number of first-shell neighbours of cell *i* (first shell defined in the Neighbour Interaction Cutoff Definitions subsection). Cell *i* thus exerts a self-propulsion force along its instantaneous polarity direction 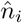:

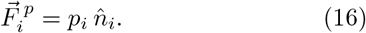

The polarity direction is updated at each timestep by blending three contributions: persistence (memory of the previous velocity direction), velocity alignment with neighbouring cells (via 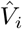, Fig. M1a), and chemosensing via the local open-area vector 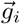 weighted by the gradient signal *χ*(*y*_*i*_) (Fig. M1b; Fig. 2A) and the polarity state *s*_*i*_ ∈ *{*+1, −1*}*:

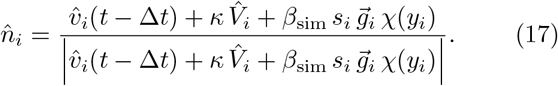

Here *κ* is the velocity-alignment coupling weight (strength of the Vicsek-like alignment relative to persistence), set to *κ* = 1 in all simulations.

The factor *s*_*i*_ implements the polarity switch: *s*_*i*_ = +1 orients propulsion *up* the gradient (chemoattraction), while *s*_*i*_ = − 1 flips the bias to orient propulsion *down* the gradient (chemorepulsion). The switching rule for *s*_*i*_ is described in the Polarity Switching Rule subsection below and analysed in detail in Section S3.

The alignment vector 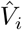 is the unit mean-velocity of the first-neighbour shell [22]:

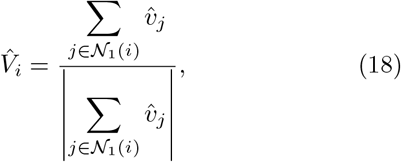

where 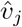 is the unit velocity of first-neighbour *j* and *N*_1_(*i*) is the first-neighbour shell of cell *i* (all cells within distance *σ*_*ij*_ + Δ_1_, see Neighbour Interaction Cutoff Definitions subsection).

### Local Open-Area Vector 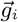

The **local open-area vector** 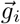 [22] (see Fig. M1b) encodes the direction and magnitude of the locally available (unoccluded) membrane of cell *i*:

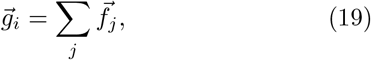

where 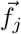 is a vector pointing in the direction bisecting the angle subtended by the centers of the *j*-th adjacent neighbor pair at the center of cell *i*, with magnitude equal to the open arc length between those two neighbors. The sum runs over all adjacent neighbor pairs of cell *i*. A core cell uniformly surrounded by neighbors has 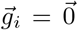 (no net open area). A rim cell with a large unoccluded arc has 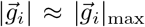, where 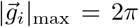 is the maximum value attained by an isolated cell with no neighbors (full circumference exposed). This captures the observation that cells polarize toward available open space: cell-cell contact blocks receptors in the contact region from ligand binding, while a larger open area allows more receptor engagement and directional sensing.

**FIG. M1.**
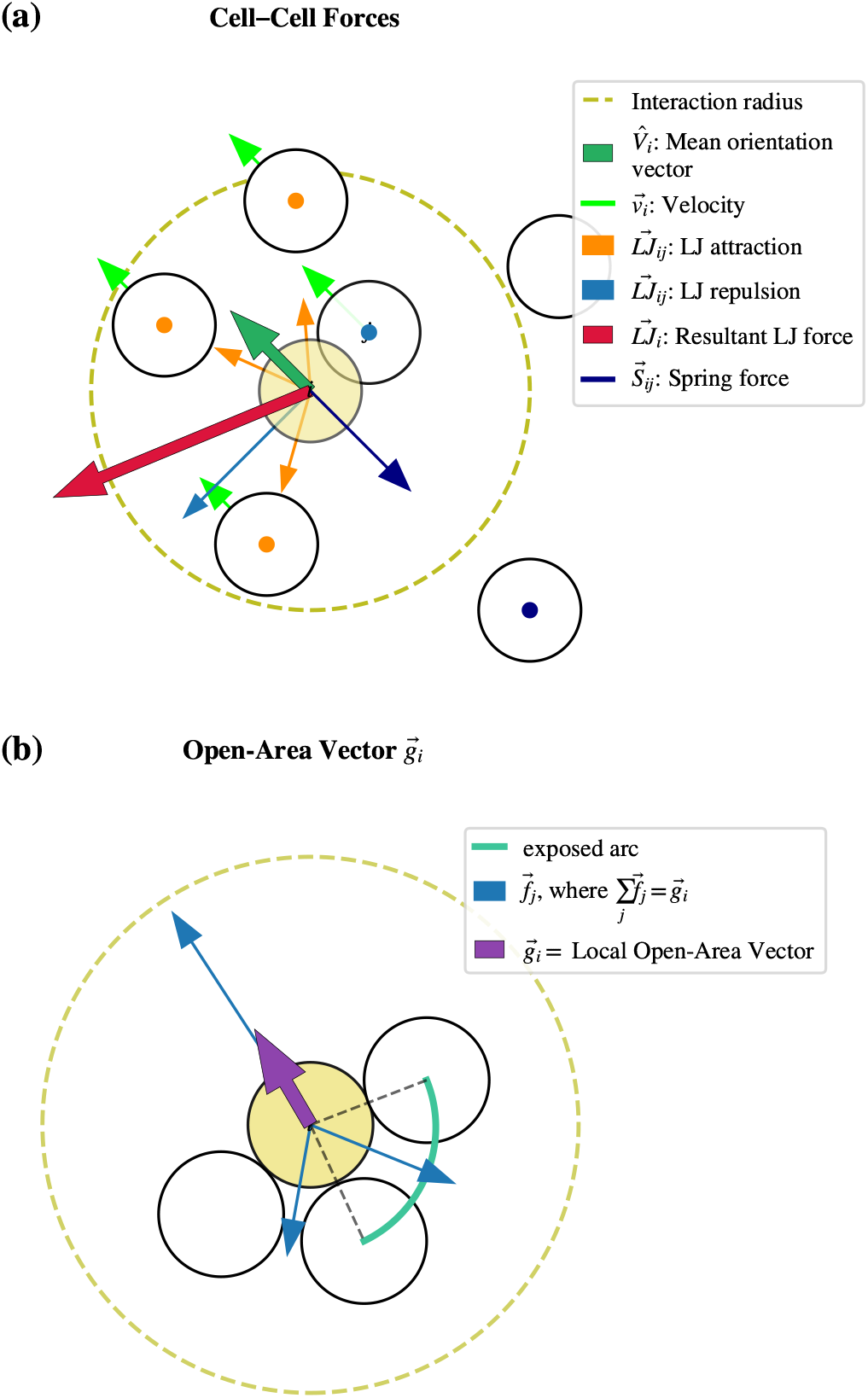
Schematic representation of the model. **(a) Cell– Cell Forces:** Illustration of the forces acting on a representative cell *i* (yellow-highlighted particle) due to its neighbors within the interaction radius (olive dashed circle). Each cell’s velocity vector 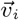 [Eq. (14)] is shown in lime green and the mean orientation vector 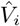[Eq. (18)] in dark green. LennardJones pairwise forces 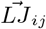 are represented as repulsive (blue arrows) for overlapping particles and attractive (orange arrows) for non-overlapping particles, generating a resultant force 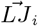 (crimson arrow) [Eq. (23)] that governs cell positioning. Spring forces 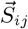 (navy arrow) [Eq. (24)] act between particles separated beyond the LJ range but within a second-neighbor cutoff, provided no intermediate particle lies between them. **(b) Local Open-Area Vector** 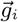: For cell *i* (yellow-highlighted), each adjacent pair of neighboring cells defines an open boundary arc (green) at the periphery of the interaction radius. The corresponding component vector 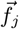 (blue arrows) points along the bisector of each arc, with magnitude proportional to the arc length. Summing all such 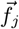 yields the local open-area vector 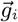 (purple arrow) [Eq. (19)], which encodes the direction of least crowding around cell *i*. The polarity switching rule is illustrated in Fig. 2A.

### Local Sensing Variable 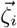

The **local sensing variable** 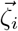 [24] combines geometric access to the external gradient with saturating receptor-ligand activation, serving as a coarse-grained proxy for effective ligand exposure of cell *i*:

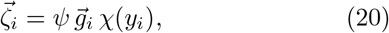

where *ψ* is the overall sensing strength. Note that 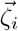 does *not* carry the polarity variable *s*_*i*_: the magnitude 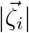 determines *whether* a cell switches (threshold criterion below), while *s*_*i*_ determines the *direction* of the chemosensing bias in (17).

**Gradient and receptor saturation**. The receptor saturation factor *χ*(*y*_*i*_) [24] is the bound fraction of CCR7 receptors at position *y*_*i*_, modelled by Langmuir adsorption:

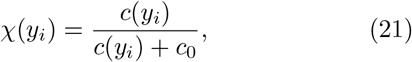

where *c*_0_ = 40 is the half-saturation constant for CCL19-CCR7 binding. The local CCL19 concentration is a linear gradient along the migration axis *y*:

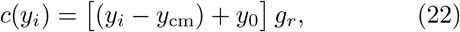

where *y*_cm_ is the cluster centre-of-mass *y*-position, *g*_*r*_ = 0.008 sim. units^−1^ is the gradient slope, and *y*_0_ = 250 sim. units ensures non-negative concentrations throughout the domain.

### Cell-Cell Contact Forces

#### Lennard-Jones (LJ) interaction

At short range, particles interact through an LJ force that captures both repulsion (preventing overlap) and attraction (maintaining cohesion) [22] (Fig. M1a):

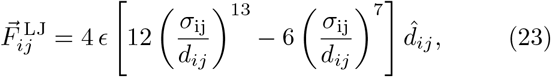

where *ϵ* is the LJ well depth, *σ*_*ij*_ = (*σ*_*i*_ +*σ*_*j*_)*/*2 is the pair contact diameter, 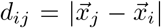 is the centre-to-centre distance, and 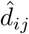 is the unit vector from *i* to *j* (Fig. M1a). The LJ force is applied within the first-neighbour shell only (Eq. 27).

#### Cohesive spring

[22] Beyond the LJ range, an additional spring force engages, providing a long-range contractile mechanism that maintains structural rigidity:

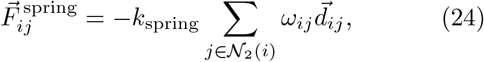

where *k*_spring_ is the spring constant, 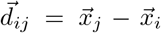 is the displacement vector, and the sum runs over second-neighbour shell *N*_2_(*i*) (see Eq. 28).

The switching variable *ω*_*ij*_ ∈ {0, 1} [22] is zero if there is an obstructing intervening particle between particles *I* and *j* (i.e., when the perpendicular distance of the intervening particle from the line joining *i* and *j* is less than *r*). If there is no such intervening particle then:

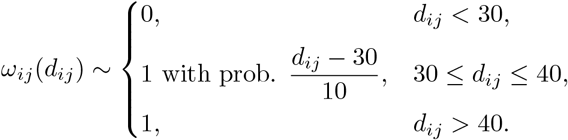

where the thresholds 30 and 40 correspond to the cell contact diameter and contact diameter plus one buffer unit respectively (in simulation units). Springs therefore activate only for unobstructed second-neighbours whose separation exceeds contact distance. The full contact force on cell *i* is

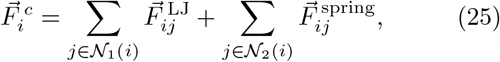

where *N*_2_(*i*) is the second-neighbour list.

### Stochastic Noise

This term captures stochastic fluctuations in actin dynamics, receptor trafficking, and protrusion activity that are not resolved at the individual cell level [22].

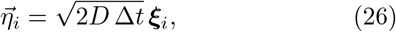

where *D* is the noise strength (diffusion coefficient), Δ*t* is the integration time step, and ***ξ***_*i*_ is a Gaussian random vector with zero mean and unit variance.

### Polarity Switching Rule

The switching criterion, Heaviside-gated rates, and polarity variable *s*_*i*_ are defined in the Model section (Eqs. 6– 10). The quantities specific to the numerical implementation are given here.

The threshold ζ_cutoff_ is a global constant calibrated to ζ_cutoff_ = 11.1 simulation units (see SI Appendix, Section S3 for the ζ_cutoff_ sensitivity sweep). For an isolated cell, all membrane is exposed so 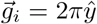 (full circumference, pointing along the external gradient), giving 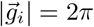, and the switching criterion reduces to a purely concentration-dependent condition. The threshold is set as ζ_cutoff_ = 0.022 ζ_max_, where ζ_max_ = *β*_sim_ · 2*π* · *χ*(*c*_max_) is the maximum sensing variable of an isolated cell at the upper boundary of the gradient.

The calibrated rates in simulation units are 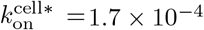 per step and 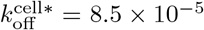 per step (Table I), corresponding to 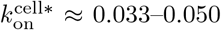 and 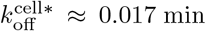 in physical units, consistent with measured CCR7 internalization and recycling timescales [39, 41].

**TABLE I.**
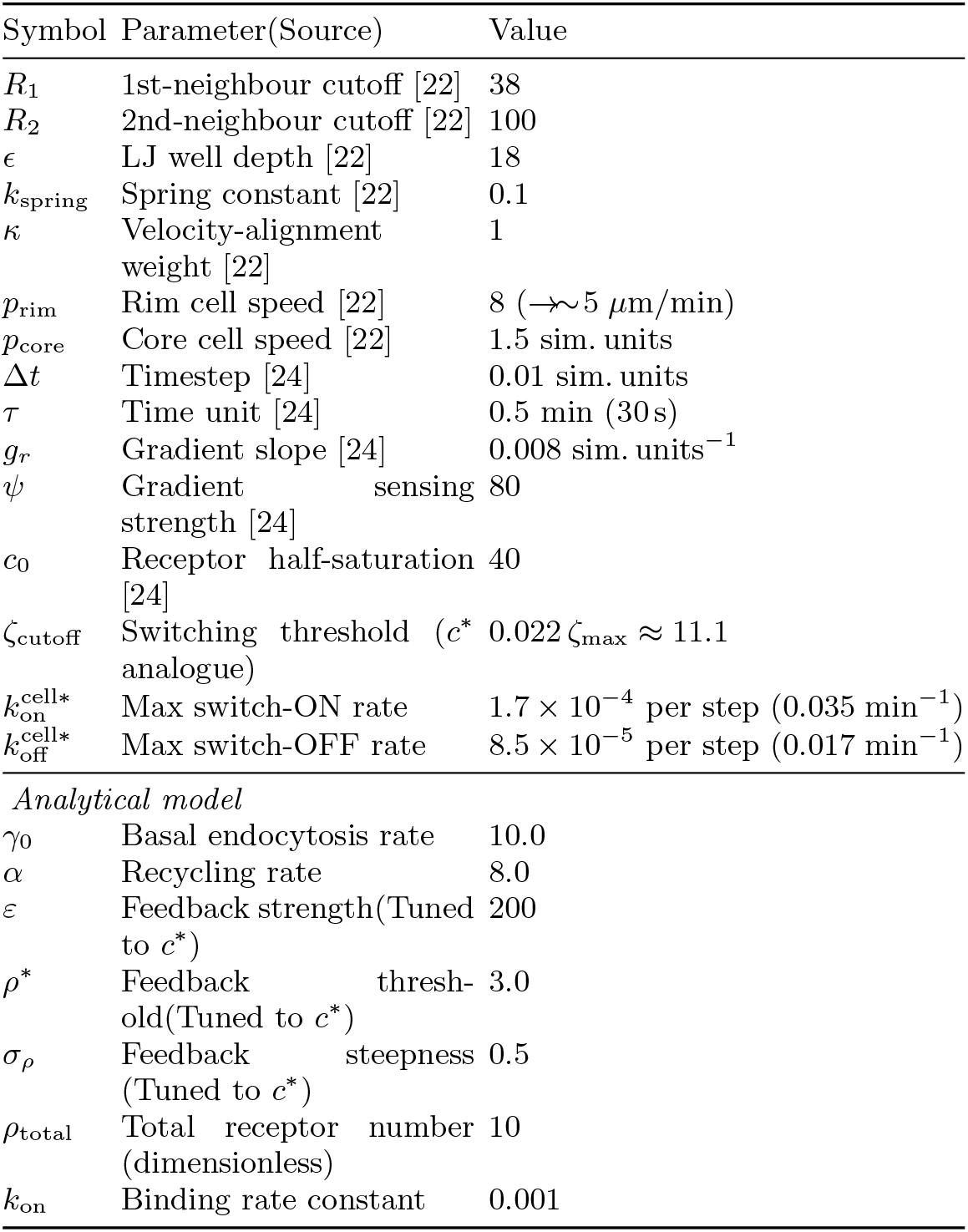
Model parameter values and sources.

### Neighbour Interaction Cutoff Definitions

Interactions are assigned according to two concentric neighbour shells centred on each particle. The **first shell** captures cells close enough to exert LJ forces. Because the system is polydisperse, the cutoff is referenced to the pair contact diameter *σ*_*ij*_ = (*σ*_*i*_ + *σ*_*j*_)*/*2, where *σ*_*i*_ is the diameter of cell *i*:

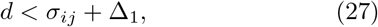

where Δ_1_ = 8 is a small buffer added beyond the contact distance (nominal diameter 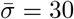, giving 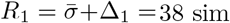. units).

The **second shell** encompasses cells outside the first shell but within reach of the cohesive spring:

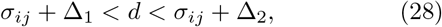

where Δ_2_ = 70 (*R*_2_ = 100 sim. units). For the nominal particle diameter, the absolute first and second cutoffs are therefore 38 and 100 sim. units, respectively [22].

### Simulation Setup and Boundary Conditions

Each simulation constrains the cluster centre-of-mass (CM) to remain stationary, so the CCL19 gradient seen by the cluster does not drift over time. This CM-fixed boundary condition is equivalent to observing the cluster in its own co-moving frame, removing net translational drift while leaving internal rearrangements, rim-core exchange, and polarity dynamics fully unconstrained. Cluster sizes from *N* = 7 to 127 cells were studied, and each run was integrated for 4,000 time steps (Δ*t* = 0.01 sim. units = 0.005 min per step), a duration sufficient for all observables to reach a statistical steady state.

### Physical Unit Conversions

Unit conversions follow Sanoria et al. [24]: one simulation step corresponds to 30 s, the nominal cell diameter of 30 sim. units maps to ≈ 15 *µ*m. Consequently, 4000 simulation time steps represent approximately 33.3 hours of experimental time.

### Parameter Calibration

Parameters are listed in Table I below, with sources indicating whether each value is directly constrained by published CCR7 kinetic data, set by cell geometry, or calibrated to satisfy cluster integrity. Only ζ_cutoff_ is fitted to the experimental FMI sign change.

## DATA AND CODE AVAILABILITY

All simulation code (Python) implementing the analytical receptor model and ABM is publicly available at https://github.com/[REPO] (DOI: [ZENODO DOI to be added upon acceptance])). Experimental data supporting the figures are from Malet-Engra et al. [31].

## ACKNOWLEDGMENTS

This work was supported by the National Science Foundation (NSF-DMS-1616926 to A.G.) and NSF-CREST: Center for Cellular and Bio-molecular Machines at UC Merced (NSF-HRD-1547848 and EES-2112675 to A.G.). M.S. acknowledge support of the Graduate Dean’s Excellence Postdoctoral Fellowship funded by Gordon and Betty Moore Foundation. A.G. and M.S. also acknowledge partial support from the NSF Center for Engineering Mechanobiology grant CMMI-154857 and computing time on the Multi-Environment Computer for Exploration and Discovery (MERCED) cluster at UC Merced (NSF-ACI-1429783). This research also benefited from the Center for Living Systems (NSF grant no. 2317138) N.S.G. is supported by the Lee and William Abramowitz Professorial Chair of Biophysics (Weizmann Institute) with additional support from a Royal Society Wolfson Visiting Fellowship. Work in G.S.’s laboratory was supported by ERC-Synergy (Grant# 801 101071470), AIRC-IG (Grant#22821), AIRC 5×1000 (#22759), the Italian Ministry of University and Research (PRIN202223GSCIT 01/ G53D23002570006/20229RM8A 001; COM-BINE/ G53D23007040001/ P2022RH4HH002; PNRR CN3RNA SPOKE/ G43C22001320007).

