## Supplementary Material for "Cell Cluster Geometry and Fluidity Control the Transition from Single-Cell Chemorepulsion to Collective Chemotaxis"

July 4, 2026

This Supplementary Information is organized as follows. Section S1 derives the steady-state receptor densities of the analytical trafficking model and provides numerical validation of the $c^*(\phi)$ scaling. Section S2 defines all simulation observables and provides multi-observable validation of the calibrated parameter set. Section S3 presents a sweep of the switching threshold $\zeta_{\mathrm{cutoff}}$ confirming that the FMI sign reversal is robust to the choice of this parameter. Section S4 shows the power-law scaling of the chemorepulsive fraction with cluster size and the collapse of FMI onto a universal critical threshold. Section S5 derives a closed-form analytical estimate of the critical repulsive fraction required to reverse collective migration. Section S6 describes the solid-cluster control experiment and its implications for cluster fluidity.

### S1. ANALYTICAL RECEPTOR TRAFFICKING MODEL

#### S1.1. Steady-State Receptor Densities

Starting from the dimensionless dynamical equations of the main text (Eqs. 1 to 3), we set the time derivatives to zero and solve for the steady-state receptor densities. The three receptor densities then satisfy:

$$\rho_{\mathrm{a}}^* = \frac{\tilde{k}_{\mathrm{on}}\,\tilde{\alpha}\,\rho_{\mathrm{total}}}{\tilde{\beta}\,\tilde{\alpha} + \tilde{k}_{\mathrm{on}}(\tilde{\beta} + \tilde{\alpha})} \tag{S1}$$

$$\rho_{\mathrm{ai}}^* = \frac{\tilde{\beta}}{\phi\,\tilde{\alpha}}\,\rho_{\mathrm{a}}^* \tag{S2}$$

$$\rho_{\mathrm{n}}^* = \rho_{\mathrm{total}} - \rho_{\mathrm{a}}^* - \phi\,\rho_{\mathrm{ai}}^* \tag{S3}$$

where $\rho_{\text{total}}$ is the conserved total receptor number (surface + endosomal), and $\tilde{\beta}$ is determined self-consistently through the intracellular feedback function:

$$\tilde{\beta} = 1 + \varepsilon \, \tfrac{1}{2} \left[ 1 + \tanh\!\left( \frac{\rho^*_{\text{ai}} - \rho^*}{\sigma_\rho} \right) \right], \tag{S4}$$

This defines a positive feedback loop. Elevated $\rho^*_{\text{ai}}$ activates the feedback signal, increasing $\tilde{\beta}$ and driving further internalization. From these solutions, the critical concentration $c^*$ (where $d\rho_a/dc = 0$, Fig. S1B) can be evaluated numerically as a function of $\phi$.

$c^*$ increases monotonically with $\phi$: cells with more cell-cell contacts (larger $\phi = V/A_{\text{exp}}$) require *higher* ligand concentrations to trigger feedback-driven chemorepulsion. Interior cluster cells with $\phi \gg 1$ have $c^*$ pushed beyond the physiological gradient window, restoring chemotaxis. For an isolated cell ($\phi = 1$), numerical solution gives $c^* = 172$ (dimensionless units), well within the physiological window (Fig. S1A). Numerical evaluation of $c^*(\phi)$ reveals an approximately linear relationship (Fig. S1C); at $\phi = 2$ (50% membrane contact) $c^* \approx 501$, and at $\phi = 4$ (75% contact) $c^*$ lies beyond the physiological gradient range.

The $\phi$ scaling enters through the internalization term in Eq. (3) of the main text: $d\rho_{\text{ai}}/d\tilde{t} = \phi^{-1}\,\tilde{\beta}\,\rho_{\text{a}} - \tilde{\alpha}\,\rho_{\text{ai}}$. Larger $\phi$ reduces the internalization flux per unit endosomal volume, preventing $\rho_{\text{ai}}$ from accumulating above $\rho^*$ except at higher $c$. Simultaneously, the recycling flux to the surface ($\phi\,\tilde{\alpha}\,\rho_{\text{ai}}$ in Eq. 2) is enhanced, replenishing $\rho_{\text{n}}$ faster and maintaining a higher surface receptor density. Together, these two effects raise the ligand threshold for chemorepulsion.

For a spherical cell of radius $R$ (normalized $R/3 = 1$):

| Cell state | Exposed fraction | $\phi$ |
|---|---|---|
| Isolated cell | 1 | 1 |
| Cluster cell, 50% contact | 0.5 | 2 |
| Cluster cell, 75% contact | 0.25 | 4 |

**Calibration of $\zeta_{\text{cutoff}}$ from the isolated-cell analytical result.** Fig. S1D shows that $\zeta(c^*(\phi)) = \psi\,|\vec{g}_i|\,\chi(c^*)$ is approximately constant across all physiological $\phi$ values (mean $\approx 0.37 \pm 0.02$, blue band). The isolated-cell reference point ($\phi = 1$, $c^* = 172$, red star) anchors this near-constant value. In the ABM, the maximum sensing variable attainable by an isolated cell (full membrane exposed, $|\vec{g}_i| = 2\pi$, at the upper gradient boundary $c_{\max}$) is

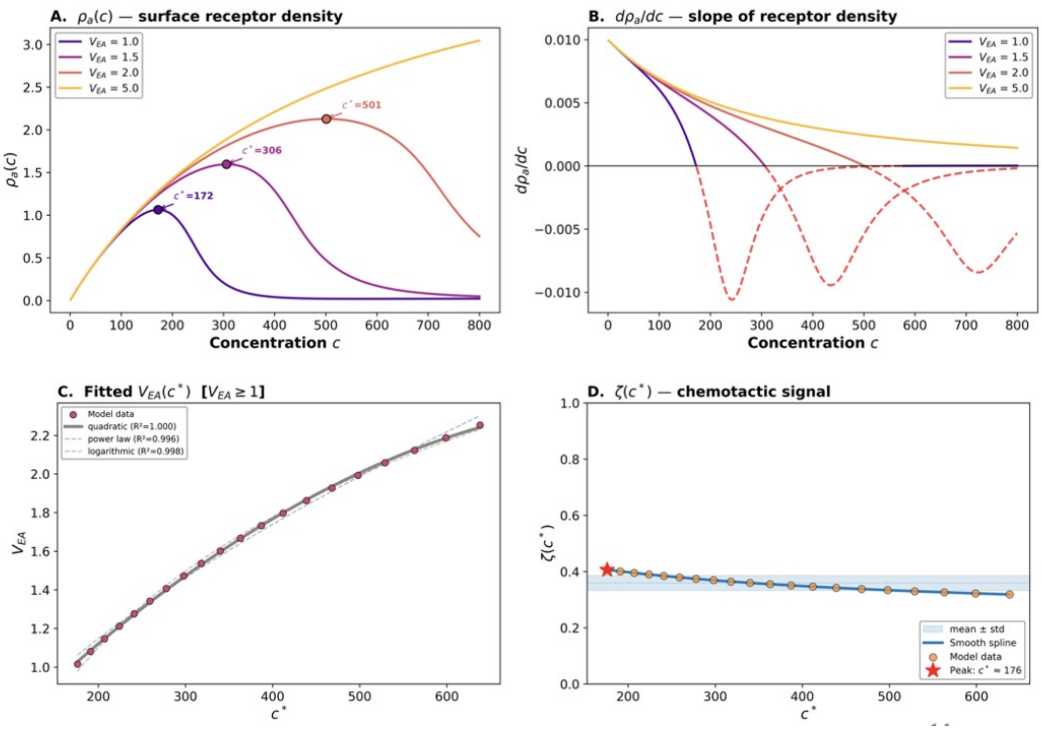


FIG. S1. **Analytical model: receptor density curves, critical concentration scaling, and self-consistency of $\zeta_{\text{cutoff}}$.** **(A)** Steady-state activated surface receptor density $\rho_a(c)$ as a function of CCL19 concentration $c$, computed numerically for $\phi = 1.0$, 1.5, 2.0, and 5.0. Each curve peaks at the critical concentration $c^*$ (filled circles), above which $d\rho_a/dc < 0$ and the cell enters the chemorepulsive regime. At $\phi = 5.0$ no peak appears within 0 to 800 ng/mL, consistent with the abolition of single-cell chemorepulsion in large clusters. **(B)** Slope $d\rho_a/dc$ of the receptor density curves in (A). Dashed portions mark regions where the slope is negative; these regions correspond to the chemorepulsive concentration window for each $\phi$. **(C)** Volume-to-exposed-area ratio $\phi$ as a function of $c^*$ (model data, pink circles). Three functional fits are shown: quadratic ($R^2 = 1.000$), power law ($R^2 = 0.996$), and logarithmic ($R^2 = 0.998$). The quadratic fit is essentially exact, indicating that $\phi$ increases faster than linearly with $c^*$ across the experimentally relevant range. The simple proportionality $c^* \propto \phi$ (scaling derived numerically) is an analytical approximation valid near $\phi \approx 1$; the full numerical relationship is captured by the quadratic fit. **(D)** The ABM sensing variable evaluated at the polarity inversion point, $\zeta(c^*) = \beta_{\text{sim}} |\vec{g}_i| \chi(c^*)$, plotted against $c^*$. The red star marks the isolated-cell value ($c^* = 172$, $\phi = 1$); orange circles span the cluster-cell range. Despite the nonlinear Langmuir saturation ($c_0 \approx 250$ ng/mL in analytical model units, so $c^*/c_0 \approx 0.41$ to 2.6 across the range), $\zeta(c^*)$ varies by <20% (mean $\approx 0.37 \pm 0.02$, blue band), confirming that a single universal threshold $\zeta_{\text{cutoff}}$ is a valid approximation for all $\phi$ values encountered in the ABM.

$\zeta_{\max} = \psi \cdot 2\pi \cdot \chi(c_{\max})$. The threshold is therefore the value of $\zeta$ at which an isolated cell reaches polarity inversion:

$$\zeta_{\text{cutoff}} = \zeta(c^*(\phi{=}1)) = 0.022\,\zeta_{\max} \approx 11.1 \text{ simulation units.} \tag{S5}$$

Because $\zeta(c^*(\phi))$ is approximately constant across $\phi$ (Fig. S1D), this single threshold applies to all cluster sizes without re-fitting. Robustness to the precise value of $\zeta_{\text{cutoff}}$ is demonstrated in Section S3.

### S2. ABM CALIBRATION

#### S2.1. Simulation-to-Experiment Concentration Mapping

In the simulation, the chemorepulsive ligand field is parameterised by two quantities: the gradient steepness $g_r$ and the background concentration $y_0$. The local ligand concentration experienced at the cluster centre of mass is given by $\langle c \rangle = y_0\, g_r$ (simulation units). The gradient steepness $g_r$ is calibrated against experimental CCL19 gradients: in our previous work [1], $g_r = 0.006$ was shown to correspond to a 0 to 100 ng/mL CCL19 gradient, while in the present study $g_r = 0.008$ corresponds to a 0 to 500 ng/mL gradient. Varying $y_0$ at fixed $g_r = 0.008$ therefore sweeps the average background CCL19 concentration at the cluster centre from low to high, playing the same role as the average CCL19 slab concentration in the experiment (Figs. 3E,F and 4D,E of the main text). This gradient-steepness calibration provides the physical concentration scale; the background level $y_0$ then serves as the simulation analogue of the experimental CCL19 slab concentration, with no additional free parameters.

#### S2.2. Observable Definitions

**Forward Migration Index (FMI).** The FMI quantifies directional persistence of cluster migration along the chemokine gradient axis ($y$):

$$\text{FMI} = \frac{\sum_t \left[ y_{\text{COM}}(t+1) - y_{\text{COM}}(t) \right]}{\sum_t \left| \vec{r}_{\text{COM}}(t+1) - \vec{r}_{\text{COM}}(t) \right|} \tag{S6}$$

The sum runs over all saved frames after equilibration. FMI $\in [-1, +1]$: positive values indicate migration toward the high-concentration source (chemotaxis); negative values indicate migration away (chemorepulsion). Values are reported as mean $\pm$ SEM across all independent simulation runs.

**Rim-to-Core Exchange Time.** The exchange time $\tau_{\text{exchange}}$ is the mean time between successive rim-to-core transitions, averaged across all cells and runs. Each cell $i$ carries a rim/core label $\sigma_i \in \{0, 1\}$ updated at every saved frame: $\sigma_i = 1$ if the cell is at the rim ($|\vec{g}_i| > |\vec{g}|_{\text{threshold}}$), $\sigma_i = 0$ otherwise. A rim-to-core transition event for cell $i$ is recorded when $\sigma_i$ changes from 1 to 0 between consecutive saved frames.

**Leader Cell Dwell Time.** The leader cell dwell time $\tau_{\text{leader}}$ is the mean duration a cell spends continuously at the leading-edge rim, averaged across all cells and runs at each $N$. A leader cell is defined as any rim cell ($\sigma_i = 1$) whose $y$-position exceeds the cluster centre-of-mass: $y_i > y_{\text{cm}}$ (front half of the cluster, facing the gradient source). A leader dwell event begins when a cell first satisfies both conditions and ends when it moves to the core ($\sigma_i = 0$) or to the rear rim ($y_i \leq y_{\text{cm}}$).

**Center-of-Mass Migration Speed.** The mean center-of-mass migration speed is computed as the total path length divided by total simulation time:

$$v_{\text{CM}} = \frac{1}{T} \sum_t \left| \vec{r}_{\text{COM}}(t+1) - \vec{r}_{\text{COM}}(t) \right| / \Delta t_{\text{phys}} \tag{S7}$$

where $T$ is the number of saved frames after equilibration and $\Delta t_{\text{phys}} = 0.005$ min is the physical timestep. Values are reported as mean $\pm$ SEM across runs.

**Chemorepulsive Fraction.** The chemorepulsive fraction $f$ is the instantaneous fraction of all cells in the cluster in the repulsive polarity state:

$$f = \frac{N_{\text{rep}}}{N} \tag{S8}$$

where $N_{\text{rep}}$ is the number of cells with $s_i = -1$ and $N$ is the total number of cells in the cluster at a given frame. The time-averaged $\langle f \rangle$ is used in Fig. 4B,E of the main text.

#### S2.3. Validation of Additional Observables

To confirm that the calibrated parameter set reproduces experimental dynamics beyond FMI, we measure two additional observables from the ABM (both defined in Section S2 S2.2):

the rim-to-core exchange time $\tau_{\text{exchange}}$ and the leader cell dwell time $\tau_{\text{leader}}$. Exchange time is reported for $N = 9$ to 127 at the standard gradient ($\nabla c = 0.008$, mean $\pm$ SEM, $n \geq 10$ runs per condition). Leader dwell time is reported for $N = 9$ to 61 at the calibrated threshold ($\zeta_{\text{cutoff}} = 0.022\,\zeta_{\max} \approx 11.1$, mean $\pm$ SEM, $n \geq 10$ runs per condition). Both observables are plotted as a function of cluster size $N$ in Fig. S2. The experimentally reported range is 8 to 15 min [2]. At the biologically relevant cluster size ($N \approx 9$), both timescales are consistent with the experimentally reported range ($\tau_{\text{ex}} \approx 14$ min; $\tau_{\text{leader}} \approx 12$ min). For larger clusters, both timescales decrease monotonically, reflecting the increasing frequency of rim-core exchanges as the rim population grows with $N$.

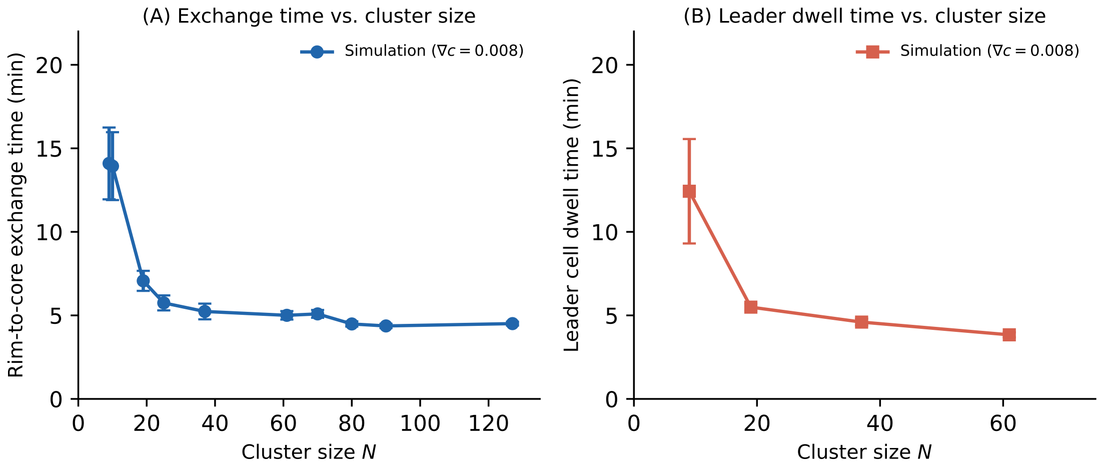


FIG. S2. **Temporal validation: rim-to-core exchange time and leader cell dwell time vs. cluster size. (A)** Mean rim-to-core exchange time $\tau_{\text{ex}}$ as a function of cluster size $N$ ($N = 9$ to 127, mean $\pm$ SEM). The experimentally reported range is 8 to 15 min [2]. At the experimentally relevant cluster size ($N \approx 9$), $\tau_{\text{ex}} \approx 14$ min is consistent with this range; larger clusters show faster exchange due to the growing rim population. **(B)** Mean leader cell dwell time $\tau_{\text{leader}}$ vs. $N$ at the calibrated cutoff ($\zeta_{\text{cutoff}} = 0.022\,\zeta_{\max} \approx 11.1$, $N = 9$ to 61, mean $\pm$ SEM). At $N = 9$, $\tau_{\text{leader}} \approx 12$ min is within the experimentally reported range; the dwell time decreases with cluster size as more rim cells compete for the leading position. Together, panels (A) and (B) show that the calibrated parameter set quantitatively reproduces experimentally measured timescales at the relevant cluster size, while predicting how these dynamics scale with cluster size.

**CCR7 surface fluctuations and rim-core exchange.** The leader cell dwell time $\tau_{\text{leader}}$ provides a mechanistic interpretation of the periodic CCR7 surface receptor fluctuations observed experimentally in leading-edge cells [2]. During prolonged residence at the rim, a leader cell is continuously exposed to high local CCL19. Feedback-driven endocytosis (Eq. S4) progressively depletes surface CCR7 over the timescale $\tau_{\text{leader}}$, reducing $\rho_{\text{a}}$ and eventually switching the cell to the chemorepulsive state. When the cell is displaced inward to the core, exposure to high CCL19 ceases, feedback weakens, and surface CCR7 is replenished through recycling (rate $\tilde{\alpha}$) over the exchange timescale $\tau_{\text{exchange}}$. Each complete depletion-recovery cycle therefore corresponds to one rim-core exchange event, and the predicted fluctuation period is $\tau_{\text{leader}} + \tau_{\text{exchange}}$. At $N \approx 9$, the model gives $\tau_{\text{leader}} \approx 12\,\text{min}$ and $\tau_{\text{exchange}} \approx 14\,\text{min}$ (Fig. S2), yielding a cycle period of $\approx 26\,\text{min}$, consistent with the experimentally reported CCR7 surface fluctuation timescale [2].

#### S2.4. Center-of-Mass Migration Speed

Center-of-mass migration speed $v_{\text{CM}}$ (defined in Section S2 S2.2) is measured across cluster sizes $N = 7$ to 127 at the standard gradient ($\nabla c = 0.008$, mean ± SEM, $n \geq 10$ runs per condition). The dashed line marks the experimentally reported value of $\approx 4\,\mu\text{m/min}$ [2]. Simulated speeds remain close to the experimental reference across all tested cluster sizes, consistent with migration driven by collective polarity rather than single-cell motility alone (Fig. S3).

### S3. $\zeta_{\text{cutoff}}$ SENSITIVITY SWEEP

The switching threshold $\zeta_{\text{cutoff}}$ is the ABM analogue of $c^*$: a cell switches to the chemorepulsive state when $|\vec{\zeta}_i| > \zeta_{\text{cutoff}}$. To confirm that the FMI sign reversal is not an artifact of a single parameter choice, we sweep $\zeta_{\text{cutoff}}$ across its full range for clusters of four sizes (Fig. S4).

$\zeta_{\text{crit}}$ **dependence on cluster size.** The value of $\zeta_{\text{cutoff}}$ at which FMI = 0 decreases monotonically with $N$ (Fig. S4B), reflecting the geometry of cluster shielding: as $N$ increases, the core fraction grows, core cells remain locked in the chemoattractive state, and the cluster-level effective switching threshold decreases. The calibrated value $\zeta_{\text{cutoff}} = 0.022\,\zeta_{\max} \approx 11.1$ lies within the sign-reversing band across all tested cluster sizes, confirming that the sign reversal is robust and not fine-tuned to a specific parameter choice. The scaling of the

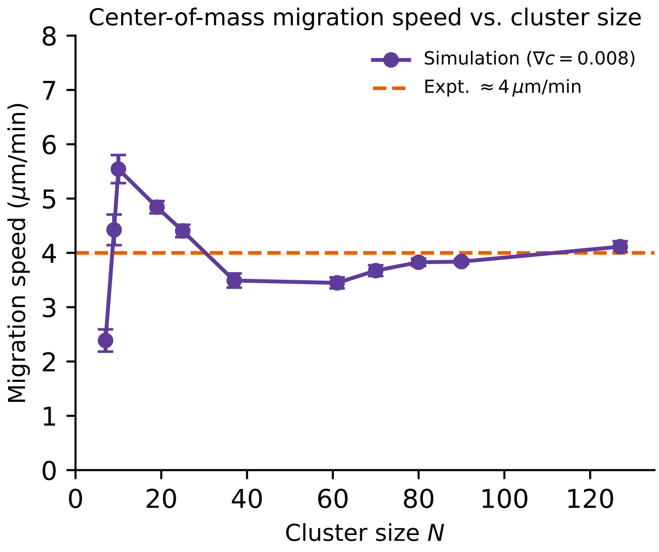


FIG. S3. **Center-of-mass migration speed vs. cluster size.** Mean $v_{\mathrm{CM}}\pm$ SEM as a function of cluster size $N$ ($N = 7$ to 127, $\nabla c = 0.008$, $n \geq 10$ runs). Dashed orange line: experimentally reported migration speed of $\approx 4\,\mu$m/min [2]. Simulated speeds are consistent with the experimental value across the full range of cluster sizes, confirming that the calibrated parameter set reproduces bulk migration dynamics without per-condition tuning.

chemorepulsive fraction $f$ with cluster size is examined in Section S4.

### S4. CHEMOREPULSIVE FRACTION SCALING AND CRITICAL SWITCHING THRESHOLD

The chemorepulsive fraction $f$ (the fraction of all cells in the cluster in the repulsive polarity state) is the key observable linking cluster geometry to collective migration direction: $f$ decays with $N$ because larger clusters have proportionally fewer rim-exposed cells, and FMI is a universal monotonically decreasing function of $f$ with a single critical threshold $f^* \approx 0.10$ that holds across all cluster sizes and gradient conditions (Fig. S5). The analytical origin of $f^*$ is derived in Section S5.

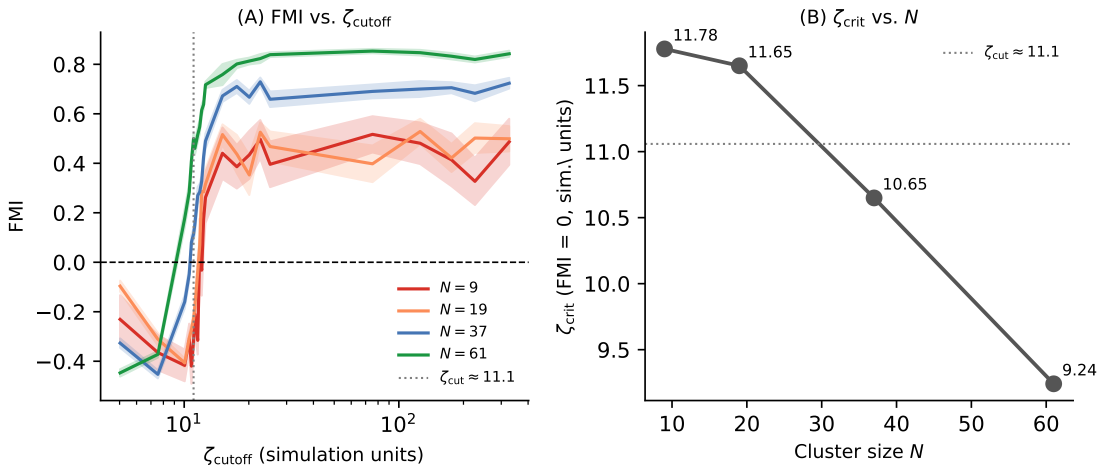


FIG. S4. **$\zeta_{\text{cutoff}}$ sweep and $\zeta_{\text{crit}}$ vs. $N$.** **(A)** Mean FMI± SEM vs. $\zeta_{\text{cutoff}}$ for cluster sizes $N \in \{9, 19, 37, 61\}$. Shaded band: range of $\zeta_{\text{cutoff}}$ over which FMI changes sign for different $N$. **(B)** $\zeta_{\text{crit}}$ (value where FMI $= 0$) vs. $N$, showing monotonic decrease (11.78, 11.65, 10.65, 9.24) as cluster size increases. Dotted line: calibrated $\zeta_{\text{cutoff}} = 0.022\,\zeta_{\max} \approx 11.1$ used in all main-text simulations.

### S5. FORCE-BALANCE ESTIMATE OF THE CRITICAL CHEMOREPULSIVE FRACTION

We derive a closed-form estimate of the critical fraction $f^*$ of chemorepulsive cells required to reverse collective migration (Fig. S6). We consider a two-layer rectangular cluster in a concentration gradient, where front cells experience higher concentration than back cells ($\lambda = F_f/F_b > 1$). Back-layer cells always exert a backward traction: in the ABM, a cell fully enclosed on its high-$c$ side has its open-area vector $\vec{g}_i$ pointing backward (into the low-$c$ space), and this toy model abstracts that geometry. Front-layer cells can be either attractive or repulsive depending on whether $c_f < c^*$ or $c_f > c^*$.

Here $n$ denotes the number of cells per layer (total cells $= 2n$) and $k$ is the number of repulsive cells in the front layer. The repulsive fraction is $f = k/(2n)$. Assuming a linear force-concentration relation $F = \mu c$, the net cluster velocity is

$$V_{\text{net}}(k) = (n-k)F_f - kF_f - nF_b = n(F_f - F_b) - 2kF_f. \tag{S9}$$

**Critical Fraction.**

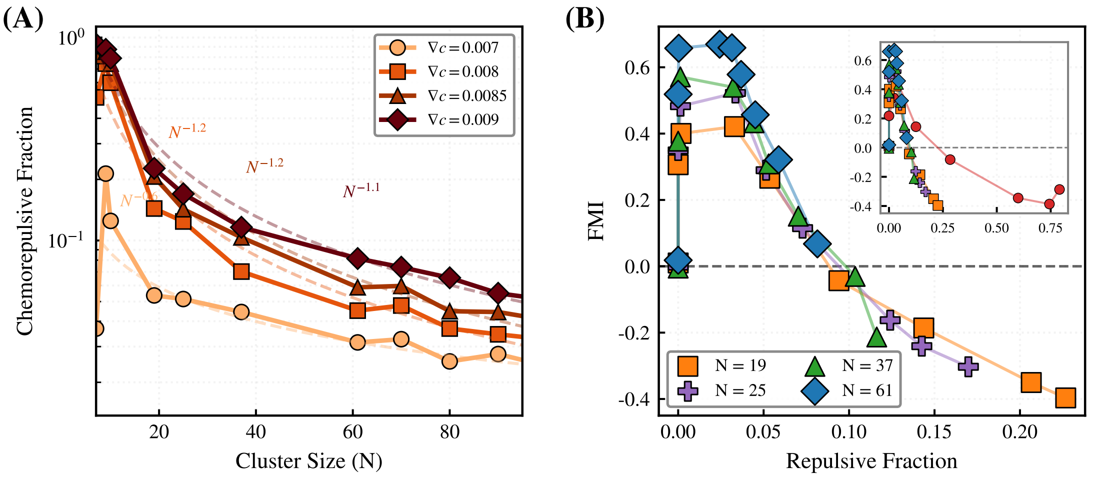


FIG. S5. **Chemorepulsive fraction scaling and critical switching threshold. (A)** Log-log plot of chemorepulsive fraction vs. $N$ for $\nabla c = 0.007, 0.008, 0.0085, 0.009$. Curves decay as $\sim N^{-0.6}$ (shallowest) to $\sim N^{-1.2}$ (steepest), with steeper gradients exceeding the $N^{-0.8}$ geometric perimeter scaling. Prefactor increases with $\nabla c$; exponent is gradient-independent, reflecting geometric control by the rim-to-total cell ratio. **(B)** FMI vs. instantaneous chemorepulsive fraction for $N = 19, 25, 37, 61$ across varying $\nabla c$. All sizes collapse onto a single monotonically decreasing curve; dashed line marks the critical threshold $f^* \approx 0.10$ separating chemoattraction (FMI $> 0$) from chemorepulsion (FMI $< 0$). *Inset*: extended to $N = 10$, confirming the collapse and threshold hold across all sizes.

Setting $V_{\text{net}} = 0$ gives

$$k^* = \frac{n(\lambda - 1)}{2\lambda}, \qquad f^* = \frac{k^*}{2n} = \frac{\lambda - 1}{4\lambda}, \tag{S10}$$

where $\lambda = F_f/F_b = c_f/c_b > 1$.

Back-layer cells always pull backward, so forward motion arises solely from attractive front cells. Migration reversal occurs when repulsive front cells cancel this forward drive. Because both attractive and repulsive front contributions scale with $F_f$, the critical fraction is bounded: $f^* < \frac{1}{4}$ for all finite $\lambda$, confirming that a minority of repulsive cells always suffices to reverse collective migration.

For example, at $\lambda = 3$, the transition occurs at $f^* = 1/6 \approx 0.17$ (Fig. S7). This analytical bound is consistent with the computationally derived threshold $f^* \approx 0.10$ from the ABM (Fig. 4D of the main text), which lies well within the upper bound. The role of cluster

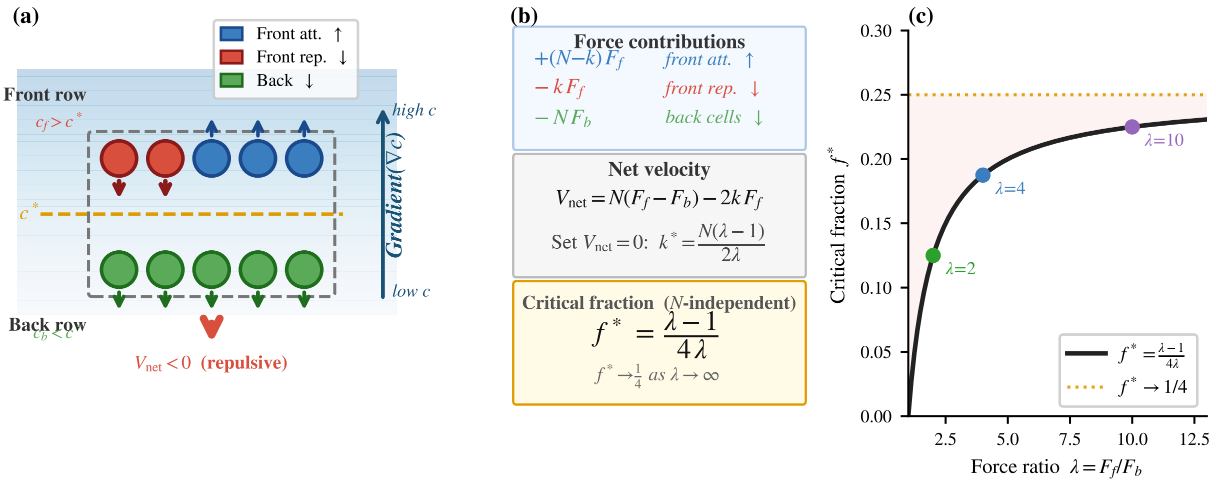


FIG. S6. **Two-layer rectangular cluster toy model. (a)** Physical setup. The cluster sits in a concentration gradient (blue shade, increasing upward). The front layer (top row, $c_f > c^*$) contains repulsive cells (red, backward ↓) and attractive cells (blue, forward ↑). The **back layer** (bottom row, $c_b < c^*$, green) always exerts a backward force $F_b$ because the only free protrusion direction is into the low-$c$ space behind the cluster. For the example shown ($k = 2$ repulsive cells, $\lambda = 3$), the net cluster velocity is backward. **(b)** Mathematical summary of the force model, net velocity equation, and the critical fraction formula.

fluidity in enabling this switching behaviour is examined in Section S6.

### S6. SOLID-CLUSTER CONTROL: INITIALIZATION AND COMPARISON

To isolate the role of cluster fluidity, we compare the full ABM against a solid-cluster variant in which internal positional exchange is suppressed, so the initial geometry is preserved throughout the simulation. This tests whether the direction switch requires dynamic rearrangement or is already encoded in geometry alone.

#### S6.1. Solid Ellipse Initialization

The solid cluster is constructed as four concentric elliptical layers with cells distributed uniformly along elliptical arcs (layer counts outer → inner: $15, 16, 16, 15$; total $N = 62$). This symmetric family scales consistently across sizes: $(2, 3, 3, 2) = 10$, $(5, 6, 6, 5) = 22$,

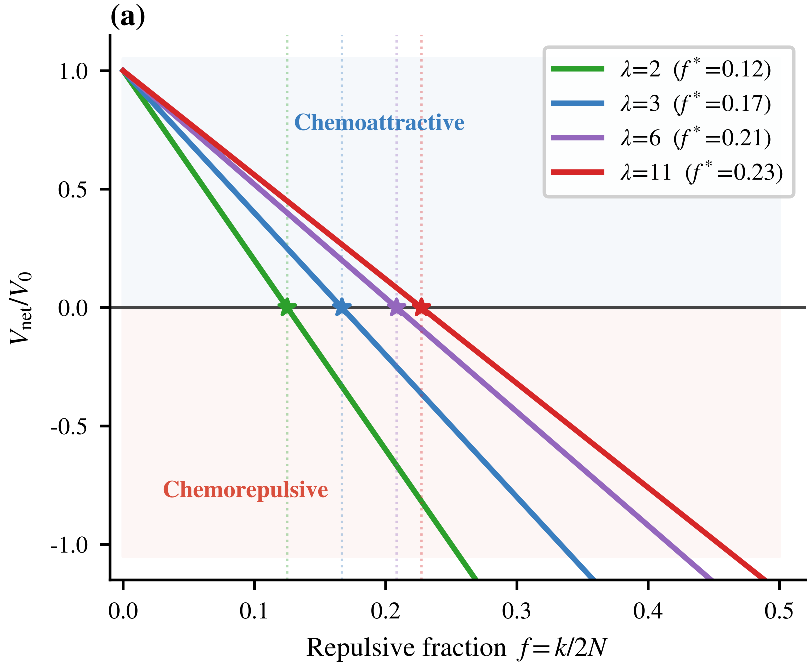


FIG. S7. **Net cluster velocity $V_{\text{net}}$ vs. chemorepulsive fraction $f$ for varying force asymmetry $\lambda = F_f/F_b$.** Net cluster velocity $V_{\text{net}}$ vs. repulsive fraction $f$ for four values of $\lambda$ (near-continuous limit, $n = 1000$); star markers show the zero-crossing $f^*$, which increases with $\lambda$ and saturates below $1/4$.

$(15, 16, 16, 15) = 62$. The ellipse aspect ratio is not pre-specified; it emerges from the layer-by-layer arc-packing procedure.

**Spring network.** Inter-cell springs are placed between all first-neighbour pairs (same distance cutoff $\bar{r} + R_1$ as the fluid model; Neighbour Interaction Cutoff Definitions, Materials and Methods). Critically, the line-of-sight (LOS) shadowing that prunes spring connections in the fluid model is *disabled*: every first-neighbour pair is spring-connected regardless of intervening cells. This denser spring network locks the relative positions of cells, suppressing positional exchange and maintaining the elliptical geometry throughout the simulation.

**Switching rates.** The off-rate $k_{\text{off}}^{\text{cell}*} = 8.5 \times 10^{-5}$ per step is the calibrated value from the fluid model (see Methods, main text).

The on-rate $k_{\text{on}}^{\text{cell}*}$ is *not* independently re-calibrated. Instead, the mean per-step transition frequency (the fraction of time a rim cell switches to the chemorepulsive state $s_i = -1$) is measured directly from fluid-cluster simulations at the same cluster size $N$, and this value is imposed as $k_{\text{on}}^{\text{cell}*}$ for the corresponding solid variant. This controlled substitution ensures that both variants have *identical switching statistics*; the only difference is whether cells can physically exchange rim and core positions.

**Evolution.** Because cells cannot exchange rim and core roles, rim cells that switch to $s_i = -1$ are mechanically trapped at the leading edge and the repulsive rim fraction grows monotonically to saturation, rather than fluctuating around a dynamic steady state as in the fluid model. Consequently, the solid cluster shows stronger repulsion at small $N$ (more rim cells permanently locked in the repulsive state) and lower positive FMI at large $N$ (persistent rim repulsion partially cancels the chemoattractive core drive) compared to the fluid variant.

This confirms that rim-to-core exchange (cluster fluidity) actively amplifies chemorepulsion at small $N$ and chemotaxis at large $N$, sharpening the behavioral switch at $N^*$. The solid-versus-fluid comparison is shown in Fig. 4C of the main text.

---

[1] M. Sanoria, G. Malet-Engra, G. Scita, N. Gov, and A. Gopinathan, PRX Life **4**, 013004 (2026).

[2] G. Malet-Engra, W. Yu, A. Oldani, J. Rey-Barroso, N. S. Gov, G. Scita, and L. Dupré, Current Biology **25**, 242 (2015).
